# Genetic mapping of sex and self-incompatibility determinants in the androdioecious plant *Phillyrea angustifolia*

**DOI:** 10.1101/2021.04.15.439943

**Authors:** Amélie Carré, Sophie Gallina, Sylvain Santoni, Philippe Vernet, Cécile Godé, Vincent Castric, Pierre Saumitou-Laprade

## Abstract

The diversity of mating and sexual systems in angiosperms is spectacular, but the factors driving their evolution remain poorly understood. In plants of the Oleaceae family, an unusual self-incompatibility (SI) system has been discovered recently, whereby only two distinct homomorphic SI specificities segregate stably. To understand the role of this peculiar SI system in preventing or promoting the diversity of sexual phenotypes observed across the family, an essential first step is to characterize the genetic architecture of these two traits. Here, we developed a high-density genetic map of the androdioecious shrub *P. angustifolia* based on a F1 cross between a hermaphrodite and a male parent with distinct SI genotypes. Using a double restriction-site associated digestion (ddRAD) sequencing approach, we obtained reliable genotypes for 196 offspring and their two parents at 10,388 markers. The resulting map comprises 23 linkage groups totaling 1,855.13 cM on the sex-averaged map. We found strong signals of association for the sex and SI phenotypes, that were each associated with a unique set of markers on linkage group 12 and 18 respectively, demonstrating inheritance of these traits as single, independent, mendelian factors. The *P. angustifolia* linkage map shows robust synteny to the olive tree genome overall. Two of the six markers strictly associated with SI in *P. angustifolia* have strong similarity with a recently identified 741kb chromosomal region fully linked to the SI phenotype on chromosome 18 of the olive tree genome, providing strong cross-validation support. The SI locus stands out as being markedly rearranged, while the sex locus has remained relatively more collinear between the two species. This *P. angustifolia* linkage map will be a useful resource to investigate the various ways by which the sex and SI determination systems have co-evolved in the broader phylogenetic context of the Oleaceae family.

## Introduction

Modes of sexual reproduction are strikingly diverse across angiosperms, both in terms of the proportion of autogamous *vs.* allogamous matings and in terms of the distribution of male and female sexual functions within and among individuals (Barrett 1998; Sakai and Weller 1999; Diggle *et al.* 2011). The conditions under which this diversity could arise under apparently similar ecological conditions and have evolved rapidly -sometimes even within the same family-have been a topic of intense interest in evolutionary biology (Barrett 1998). The control of self-fertilization and the delicate balance between its costs and benefits is considered to be a central force driving this diversity. Avoidance of self-fertilization is sometimes associated with observable phenotypic variations among reciprocally compatible partners. These variations can be morphological (e.g. distyly) or temporal (e.g. protandry, protogyny in the case of heterodichogamy), but in many cases the flowers show no obvious morphological or phenological variation, and self-fertilization avoidance relies on so-called “homomorphic” self-incompatibility (SI) systems. These systems are defined as the inability of fertile hermaphrodite plants to produce zygotes through self-fertilization (Lundqvist 1956; De Nettancourt 1977), and typically rely on the segregation of a finite number of recognition “specificities” whereby matings between individuals expressing cognate specificities are not successful at producing zygotes. At the genetic level, the SI specificities most commonly segregate as a single multi-allelic mendelian locus, the S locus. This locus contains at least two genes, one encoding the male determinant expressed in pollen and the other encoding the female determinant expressed in pistils, with the male specificity sometimes determined by a series of tandemly arranged paralogs (Kubo *et al.* 2015). The male and female determinants are both highly polymorphic and tightly linked, being inherited as a single non-recombining genetic unit. In cases where the molecular mechanisms controlling SI could be studied in detail, they were found to be remarkably diverse, illustrating their independent evolutionary origins across the flowering plants (Iwano and Takayama 2012). Beyond the diversity of the molecular functions employed, SI systems can also differ in their genetic architecture. In the Poaceae family for example, two independent loci (named S and Z) control SI (Yang, et al., 2008). In other cases, the alternate allelic specificities can be determined by presence-absence variants rather than nucleotide sequence variants of a given gene, such as *e.g.* in *Primula vulgaris*, where one of the two reproductive phenotypes is hemizygous rather than heterozygous for the SI locus (Li *et al.* 2016).

In spite of this diversity of molecular mechanisms and genetic architectures, a common feature of SI phenotypes is that they are all expected to evolve under negative frequency-dependent selection, a form of natural selection favoring the long-term maintenance of high levels of allelic diversity (Wright 1939). Accordingly, large numbers of distinct SI alleles are commonly observed to segregate within natural and cultivated SI species (reviewed in Castric and Vekemans 2004). There are notable exceptions to this general rule, however, and in some species only two SI specificities seem to segregate stably. Most often in such diallelic SI systems, the two SI specificities are in perfect association with morphologically distinguishable floral phenotypes. In distylous species, for instance, two floral morphs called “pin” (L-morph) and “thrum” (S-morph) coexist (Barrett 1992; Barrett 2019). In each morph, the anthers and stigma are spatially separated within the flowers, but located at corresponding, reciprocal positions between the two morphs. Additional morphological differences exist, with S-morph flowers producing fewer but larger pollen grains than L-morph flowers (Dulberger 1992). These morphological differences are believed to enhance the selfing avoidance conferred by the SI system but also to increase both male and female fitnesses (Barrett 1990; Barrett 2002; Keller *et al.* 2014), although it is not clear which of SI or floral morphs became established in the first place (Charlesworth and Charlesworth 1979).

The Oleacea family is another intriguing exception, where a diallelic SI system was recently found to be shared across the entire family (Vernet *et al.* 2016). In this family of trees, the genera *Jasminum* (2*n* = 26), *Fontanesia* (2*n* = 26) and *Forsythia* (2*n*= 28) are all heterostylous and are therefore all expected to possess a heteromorphic diallelic SI system; in *Jasminum fruticans* self- and within-morph fertilization are unsuccessful (DommÉe *et al.* 1992). The ancestral heterostyly gave rise to species with hermaphrodite (e.g. *Ligustrum vulgare*, *Olea europaea*), androdioecious (e.g. *P. angustifolia*, *Fraxinus ornus*), polygamous (e.g. *Fraxinus excelsior*) and even dioecious (e.g. *Fraxinus chinensis*) sexual systems, possibly in association with a doubling of the number of chromosomes (2*n*= 46 in the Oleeae tribe) (Taylor 1945; Wallander and Albert 2000). Evaluation of pollen germination success in controlled *in vitro* crossing experiments (whereby fluorescence microscopy is used to score the growth of pollen tubes reaching the style through the stigma; referred to below as the “stigma test”) revealed the existence of a previously unsuspected homomorphic diallelic SI in one of these species, *P. angustifolia* (Saumitou-Laprade *et al.* 2010). In this androdioecious species (i.e. in which male and hermaphrodite individuals coexist in the same populations), hermaphrodite individuals form two morphologically indistinguishable groups of SI specificities that are reciprocally compatible but incompatible within groups, whereas males show compatibility with hermaphrodites of both groups (Saumitou-Laprade *et al.* 2010). This “universal” compatibility of males offsets the reproductive disadvantage they suffer from lack of their female function, such that the existence of the diallelic SI system provides a powerful explanation to the long-standing evolutionary puzzle represented by the maintenance of high frequencies of males in this species (Pannell and Korbecka 2010; Saumitou-Laprade *et al.* 2010; Billiard *et al.* 2015; Pannell and Voillemot 2015). Extension of the stigma test developed in *P. angustifolia* to other species of the same tribe including *L. vulgaris* (De Cauwer *et al.* 2020), *F. ornus* (Vernet *et al.* 2016) and *O. europaea* (Saumitou-Laprade *et al.* 2017; Dupin *et al.* 2020), demonstrated that all species exhibited some form of the diallelic SI system, but with no consistent association with floral morphology. Cross-species pollination experiments even showed that pollen from *P. angustifolia* was able to trigger a robust SI response on *O. europaea* and the more distant *F. ornus* and *F. excelsior* stigmas (the reciprocal is also true). This opens the question of whether the homomorphic diallelic SI determinants are orthologs across the Oleeae tribe, even in the face of the variety of sexual polymorphisms present in the different species. More broadly, the link between determinant of the homomorphic diallelic SI in the Oleeae tribe and those of the heteromorphic diallelic SI in the ancestral diploid, largely heterostylous species, remains to be established (Barrett 2019). Understanding the causes of the long-term maintenance of this SI system and exploring its consequences on the evolution of sexual systems in hermaphrodite, androdioecious, polygamous or dioecious species of the family represents an important goal. The case of *P. angustifolia* is particularly interesting because it is one of the rare instances where separate sexes decoupled from mating types can be studied in a single species (Charlesworth 1978).

A first step toward a better understanding of the role of the diallelic SI system in promoting the sexual diversity in Oleaceae is to characterize and compare the genetic architecture of the SI and sexual phenotypes. At this stage, however, the genomic resources for most of these non-model species remain limited. In this context, the recent sequencing efforts (Unver *et al.* 2017; JimÉnez-Ruiz *et al.* 2020) and the genetic mapping of the SI locus in a biparental population segregating for SI groups in *Olea europaea* (Mariotti *et al.* 2020) represent major breakthroughs in the search for the SI locus in Oleaceae. They have narrowed down the SI locus to an interval of 5.4cM corresponding to a region of approximately 300kb, but it is currently unknown whether the same region is controlling SI in other species. In *P. angustifolia,* based on a series of genetic analysis of progenies from controlled crosses, Billiard *et al.* (2015) proposed a fairly simple genetic model, where sex and SI are controlled by two independently segregating diallelic loci. Under this model, sex would be determined by the “M” locus at which a dominant *M* allele codes for the male phenotype (*i.e. M is a* female-sterility mutation leading e.g. to arrested development of the stigma) and a recessive *m* allele codes for the hermaphrodite phenotype. The S locus would encode the SI system and comprise a dominant allele *S2* and a recessive allele *S1.* The model thus hypothesizes that hermaphrodites are homozygous *mm* at the sex locus, and fall into two groups of SI specificities, named H_a_ and H_b_ carrying the *S1S1* and *S1S2* genotypes at the S locus, respectively (their complete genotypes would thus be *mmS1S1* and *mmS1S2* respectively). The model also hypothesizes three male genotypes (M_a_: *mMS1S1*, M_b_: *mMS1S2*, and M_c_*: mMS2S2*). In addition, Billiard *et al.* (2015) experimentally showed that, while males are compatible with all hermaphrodites, the segregation of sexual phenotypes varies according to which group of hermaphrodites they sire: the progeny of H_a_ hermaphrodites pollinated by males systematically consists of both hermaphrodites and males with a consistent but slight departure from 1:1 ratio, while that of H_b_ hermaphrodites pollinated by the very same males systematically consists of male individuals only. These segregation patterns suggests a pleiotropic effect of the *M* allele, conferring not only female sterility and universal pollen compatibility, but also a complete male-biased sex-ratio distortion when crossed with one of the two groups of hermaphrodites and a more subtle departure from 1:1 ratio when crossed with the other group of hermaphrodites (Billiard *et al.* 2015). The latter departure, however, was observed on small progeny arrays only, and its magnitude thus comes with considerable uncertainty.

In this study, we developed a high-density genetic map for the non-model tree *P. angustifolia* using a ddRAD sequencing approach and used it to address three main questions related to the evolution of its peculiar reproductive system. First, are the SI and sex phenotypes in *P. angustifolia* encoded by just two independent loci, as predicted by the most likely segregation model of Billiard *et al.* (2015)? Second, which genomic regions are associated with the SI and sex loci, and what segregation model do the SI and sex-associated loci follow (i.e. which of the males or hermaphrodites, and which of the two SI phenotypes are homozygous *vs.* heterozygous at either loci, or are these phenotypes under the control of hemizygous genomic regions?). Third, what is the level of synteny between our *P. angustifolia* genetic map and the recently published Olive tree genome (Unver *et al.* 2017; Mariotti *et al.* 2020), both globally and specifically at the SI and sex-associated loci?

## Material and Methods

### Experimental cross and cartography population

In order to get both the SI group and the sexual phenotype (males vs hermaphrodites) to segregate in a single progeny array, a single maternal and a single paternal plant were chosen among the progenies of the controlled crosses produced by Billiard *et al.*(2015). Briefly, a H_a_ maternal tree (named 01.N-25, with putative genotype mmS1S1) was chosen in the progeny of a (H_a_ × M_a_) cross. It was crossed in March 2012 to a M_b_ father (named 13.A-06, putative genotype mM S1S2) chosen in the progeny of a (H_a_ × M_c_) cross, following the protocol of Saumitou-Laprade *et al.* (2010). Both trees were maintained at the experimental garden of the “Plateforme des Terrains d’Expérience du LabEx CeMEB,” (CEFE, CNRS) in Montpellier, France. F1 seeds were collected in September 2012 and germinated in the greenhouse of the “Plateforme Serre, cultures et terrains expérimentaux,” at the University of Lille (France). Seedling paternity was verified with two highly polymorphic microsatellite markers (Vassiliadis *et al.* 2002), and 1,064 plants with confirmed paternity were installed in May 2013 on the experimental garden of the “Plateforme des Terrains d’Expérience du LabEx CeMEB,” (CEFE, CNRS) in Montpellier. Sexual phenotypes were visually determined based on the absence of stigma for 1,021 F1 individuals during their first flowering season in 2016 and 2017 (absence of stigma indicates male individuals). Twenty-one progenies did not flower and 22 died during the test period. The hermaphrodite individuals were assigned to an SI group using the stigma test previously described in Saumitou-Laprade *et al.* (2010; Saumitou-Laprade *et al.* 2017).

### DNA extraction, library preparation and sequencing

In 2015, *i.e.* the year before sexual phenotypes were determined and stigma tests were performed, 204 offspring were randomly selected for genomic library preparation and genotyping. Briefly, DNA from parents and progenies was extracted from 100 mg of frozen young leaves with the Chemagic DNA Plant Kit (Perkin Elmer Chemagen, Baesweller, DE, Part # CMG-194), according to the manufacturer’s instructions. The protocol was adapted to the use of the KingFisher Flex™ (Thermo Fisher Scientific, Waltham, MA, USA) automated DNA purification workstation. The extracted DNA was quantified using a Qubit fluorometer (Thermo Fisher Scientific, Illkirch, France). Genome complexity was reduced by double digestion restriction associated DNA sequencing (ddRAD seq) (Peterson *et al.* 2012) using two restriction enzymes: *PstI*, a rare-cutting restriction enzyme sensitive to methylation recognizing the motif CTGCA/G, and *MseI*, a common-cutting restriction enzyme (recognizing the motif T/TAA). The libraries were constructed at the INRAE – AGAP facilities (Montpellier, France). Next-generation sequencing was performed in a 150-bp paired-ends-read mode using three lanes on a HiSeq3000 sequencer (Illumina, San Diego, CA, USA) at the Get-Plage core facility (Genotoul platform, INRAE Toulouse, France).

### GBS data analysis and linkage mapping

Illumina sequences were quality filtered with the *process_radtags* program of Stacks v2.3 (Catchen *et al.* 2011) to remove low-quality base calls and adapter sequences. We followed the Rochette & Catchen protocol (Rochette and Catchen 2017) to obtain a *de novo* catalog of reference loci. Briefly, the reads were assembled and aligned with a minimum stack depth of 3 (−m=3) and at most two nucleotide differences when merging stacks into loci (−M=2). We allowed at most two nucleotide differences between loci when building the catalog (−n=2). Both parental and all offspring FASTQ files were aligned to the *de novo* catalog using Bowtie2 v2.2.6 (Langmead and Salzberg 2012), the option ‘end-to-end’ and ‘sensitive’ were used for the alignment. At this step, one .bam file was obtained per individual to construct the linkage map with Lep-MAP3 (Rastas 2017). A custom python script was used to remove SPN markers with reads coverage <5. After this step, the script calls Samtools v1.3.1 and the script *pileupParser2.awk* (limit1=5) to convert *.bam* files to the format used by Lep-MAP3. We used the *ParentCall2* module of Lep-MAP3 to select loci with reliable parental genotypes by considering genotype information on parents and offspring. The *Filtering2* module was then used to remove non-informative and distorted markers (dataTolerance = 0.0000001). The module *SeparateChromosomes2* assigned markers to linkage groups (LGs), after test, where the logarithm of odds score (LodLimit) varied from 10 to 50 in steps of 5 then from 20 to 30 in steps of 1 and the minimum number of SNP markers (sizeLimit) per linkage group from 50 to 500 in steps of 50 for each of the LodLimit. The two parameters, lodLimit = 27 and sizeLimit = 250, were chosen as the best parameters to obtain the 23 linkage groups (as expected in members of the Oleoideae subfamily; Wallander and Albert 2000). A custom python script removed loci with SNPs mapped on two or more different linkage groups. The last module *OrderMarkers2* ordered the markers within each LG. To consider the slight stochastic variation in marker distances between executions, the module was run three times on each linkage group, first separately for the meiosis that took place in each parent (sexAveraged = 0) and then averaged between the two parents (sexAveraged = 1). To produce the most likely final father and mother specific maps and a final sex-averaged maps (De-Kayne and Feulner 2018), we kept for each map the order of markers that had the highest likelihoods for each linkage group. In the end of some linkage groups, we removed from the final genetic map markers that were clearly outliers i.e. that had orders of magnitude more recombination to any marker than the typical average (Table 1). The original map is provided in Figure S1.

**Table 1.** Comparison of the sex-averaged, male and female linkage maps. The values in this table are computed without the outliers SNP markers at the extremity of the linkage groups.

| Linkage group | Number of SNPs | Number of SNPs (without outliers) | Sex-averaged map |  |  | Paternal map |  |  | Maternal map |  |  |
| --- | --- | --- | --- | --- | --- | --- | --- | --- | --- | --- | --- |
|  |  |  | LG length (cM) | SNPs/cM | average intermarker distance | LG Length (cM) | SNPs/cM | average intermarker distance | LG Length (cM) | SNPs/cM | average intermarker distance |
| 1 | 854 | 839 | 75.78 | 11.07 | 0.09 | 79.30 | 10.58 | 0.09 | 92.11 | 9.11 | 0.11 |
| 2 | 633 | 621 | 23.90 | 25.98 | 0.10 | 35.00 | 17.74 | 0.12 | 22.73 | 27.32 | 0.11 |
| 3 | 676 | 676 | 74.50 | 9.07 | 0.11 | 61.94 | 10.91 | 0.09 | 85.42 | 7.91 | 0.13 |
| 4 | 535 | 535 | 68.07 | 7.86 | 0.13 | 71.63 | 7.47 | 0.13 | 69.82 | 7.66 | 0.13 |
| 5 | 502 | 494 | 56.02 | 8.82 | 0.11 | 50.58 | 9.77 | 0.10 | 67.84 | 7.28 | 0.14 |
| 6 | 877 | 877 | 96.89 | 9.05 | 0.11 | 90.81 | 9.66 | 0.10 | 103.63 | 8.46 | 0.12 |
| 7 | 609 | 601 | 64.14 | 9.37 | 0.11 | 68.93 | 8.72 | 0.11 | 64.99 | 9.25 | 0.11 |
| 8 | 486 | 479 | 62.71 | 7.64 | 0.13 | 91.89 | 5.21 | 0.19 | 119.25 | 4.02 | 0.25 |
| 9 | 408 | 406 | 63.28 | 6.42 | 0.16 | 56.06 | 7.24 | 0.14 | 71.04 | 5.72 | 0.18 |
| 10 | 1365 | 1361 | 110.69 | 12.30 | 0.08 | 121.95 | 11.16 | 0.09 | 112.38 | 12.11 | 0.08 |
| 11 | 793 | 783 | 91.66 | 8.54 | 0.12 | 80.40 | 9.74 | 0.10 | 108.84 | 7.19 | 0.14 |
| 12 | 973 | 969 | 77.12 | 12.56 | 0.08 | 91.52 | 10.59 | 0.09 | 88.09 | 11.00 | 0.09 |
| 13 | 849 | 848 | 77.40 | 10.96 | 0.09 | 77.29 | 10.97 | 0.09 | 80.77 | 10.50 | 0.10 |
| 14 | 566 | 565 | 62.12 | 9.10 | 0.11 | 72.25 | 7.82 | 0.13 | 71.39 | 7.91 | 0.13 |
| 15 | 750 | 747 | 76.92 | 9.71 | 0.10 | 82.98 | 9.00 | 0.11 | 96.01 | 7.78 | 0.13 |
| 16 | 591 | 589 | 53.53 | 11.00 | 0.09 | 56.93 | 10.35 | 0.10 | 69.56 | 8.47 | 0.12 |
| 17 | 613 | 613 | 76.52 | 8.01 | 0.13 | 70.58 | 8.69 | 0.12 | 83.94 | 7.30 | 0.14 |
| 18 | 660 | 659 | 76.22 | 8.65 | 0.12 | 77.26 | 8.53 | 0.12 | 81.99 | 8.04 | 0.12 |
| 19 | 806 | 806 | 69.29 | 11.63 | 0.09 | 91.10 | 8.85 | 0.11 | 79.49 | 10.14 | 0.10 |
| 20 | 547 | 531 | 56.26 | 9.44 | 0.11 | 67.56 | 7.86 | 0.13 | 62.91 | 8.44 | 0.12 |
| 21 | 479 | 476 | 54.86 | 8.68 | 0.12 | 69.49 | 6.85 | 0.15 | 54.14 | 8.79 | 0.11 |
| 22 | 550 | 544 | 57.05 | 9.54 | 0.11 | 55.64 | 9.78 | 0.10 | 61.44 | 8.85 | 0.11 |
| 23 | 690 | 684 | 61.64 | 11.10 | 0.09 | 67.10 | 10.19 | 0.10 | 66.42 | 10.30 | 0.10 |
| average | 687 | 682 | 68.98 | 10.28 | 0.11 | 73.40 | 9.46 | 0.11 | 78.88 | 9.29 | 0.12 |

### Sex and SI locus identification

To identify the sex-determination system in *P. angustifolia* we considered two possible genetic models. First, a “XY” male heterogametic system, where males are heterozygous or hemizygous (XY) and hermaphodites are homozygous (XX). Second, a “ZW” hermaphrodite heterogametic system, where hermaphodites are heterozygous or hemizygous (ZW) and males are homozygous (ZZ). We applied the same logic to the SI determination system, as segregation patterns (Billiard *et al.* 2015) suggested that SI possibly also has a heterogametic determination system, with homozygous H_a_ and heterozygous H_b_. In the same way as for sex, it is therefore possible to test the different models (XY, ZW or hemizygous) to determine which SNPs are linked to the two SI phenotypes.

Based on this approach, we identified sex-linked and SI-linked markers on the genetic map by employing SEX-DETector, a maximum-likelihood inference model initially designed to distinguish autosomal from sex-linked genes based on segregation patterns in a cross (Muyle *et al.* 2016). Briefly, a new alignment of reads from each individual on the loci used to construct the linkage map was done with bwa (Li and Durbin 2009). This new alignment has the advantage of retrieving more SNPs than used by LepMap3, as SNPs considered as non-informative by LepMap3 can still be informative to distinguish among sex- or SI-determination systems by SEX-DETector. The alignment was analyzed using Reads2snp (default tool for SEX-DETector) (Tsagkogeorga *et al.* 2012) with option -par 0. We ran Reads2snp without the -aeb (account for allelic expression bias) option to accomodate for the use of genomic rather than RNA-seq data. For each phenotype (H_a_ vs. H_b_ and males vs. hermaphrodites), SEX-DETector was run for both a XY and a ZW model with the following parameters: -detail, -L, -SEM, -thr 0.8, -E 0.05. For each run, SEX-DETector also calculates the probability for X (or Z)-hemizygous segregation in the heterozygous haplotypes. To compensate for the heterogeneity between the number of males (83) and hermaphrodites (113) in our progeny array, each model was tested three times with sub-samples of 83 hermaphrodites obtained by randomly drawing from the 113 individuals. We retained SNPs with a ≥80% probability of following an XY (or ZW) segregation pattern, with a minimum of 50% individuals genotyped and less than 5% of the individuals departing from this model (due to either genotyping error or crossing-over).

### Synteny analysis with the olive tree

To study synteny, we used basic local alignment search tool (BLAST) to find regions of local similarity between the *P. angustifolia* ddRADseq loci in the linkage map and the *Olea europea var. sylvestris* genome assembly (Unver *et al.* 2017). This assembly is composed of 23 main chromosomes and a series of 41,233 unanchored scaffolds for a total of 1,142,316,613 bp. Only loci with a unique hit with at least 85% identity over a minimum of 110 bp were selected for synteny analysis. Synteny relationships were visualized with *circos-0.69-6* (Krzywinski *et al.* 2009). Synteny between linkage groups of *P. angustifolia* and the main 23 *O. europea* chromosomes was established based on the number of markers with a significant BLAST hit. At a finer scale, we also examined synteny with the smaller unanchored scaffolds of the assembly, as the history of rearrangement and allo-tetraploidization is likely to have disrupted synteny.

## Results

### Phenotyping progenies for sex and SI groups

As expected, our cartography population segregated for sex and SI phenotypes, providing a powerful resource to genetically map these two traits. Among the 1,021 F1 individuals that flowered during the two seasons of phenotyping, we scored 619 hermaphrodites and 402 males, revealing a biased sex ratio in favor of hermaphrodites (khi^2^= 46.12, p-value=1.28×10^−11^). Stigma tests were successfully performed on 613 hermaphrodites (6 individuals flowered too late to be included in a stigma test), revealing 316 H_a_ and 297 H_b_, i.e. an equilibrated segregation of the two SI phenotypes (khi^2^=1.22, p-value= 0.27). The random subsample of 204 F1 progenies chosen before the first flowering season for ddRAD-seq analysis (see below) followed similar phenotypic proportions. Only 196 of the 204 progenies ended up flowering, revealing 83 males and 113 hermaphrodites, among which 60 belonged to the H_a_ group and 53 to the H_b_ group.

### Linkage mapping

The two parents and the 196 offspring that had flowered were successfully genotyped using a ddRAD-seq approach. Our stringent filtering procedure identified 11,070 loci composed of 17,096 SNP markers as being informative for Lep-MAP3. By choosing a LOD score of 27, a total of 10,388 loci composed of 15,814 SNPs were assigned to, and arranged within, 23 linkage groups in both sex-averaged and sex-specific maps (Table 1).

The linkage groups of the mother map were on average larger (78.88 cM) than the linkage groups of the father map (73.40 cM) and varied from 22.73 cM to 112.38 cM and from 35 cM to 121.94 cM respectively (Table 1, Figure S1). The total map lengths were 1586.57 cM, 1688.16 cM and 1814.19 cM in the sex-averaged, male and female maps, respectively. The length of the linkage groups varied from 23.90 cM to 110.69 cM in the sex-averaged map, with an average of 683 SNPs markers per linkage group (Table 1).

### Sex and SI locus identification

We found evidence that a region on linkage group 18 (LG18) was associated with the SI phenotypes, with Hb hermaphrodites having heterozygous genotype, akin to a XY system. Indeed, when comparing H_a_ and H_b_, among the 38,998 SNPs analyzed by SEX-DETector, 496 had a probability of following an XY pattern ≥0.80. We then applied two stringent filters by retaining only SNPs that had been genotyped for more than 50% of the offspring (n=211), and for which less than 5% of the offspring departed from the expected genotype under a XY model (n=23). Six of these 23 SNPs, distributed in 4 loci, followed a segregation pattern strictly consistent with a XY model. These four loci are tightly clustered on the linkage map and define a region of 1.230 cM on LG18 (Figure 1) in the sex-averaged map. Relaxing the stringency or our thresholds, this region also contains five loci that strictly follow an XY segregation but with fewer than 50% of offsprings successfully genotyped, as well as six loci with autosomal inheritance, possibly corresponding to polymorphisms accumulated within allelic lineages associated with either of the alternate SI specificities. Using the same filtering scheme, none of the SNPs was found to follow a ZW pattern.

**Figure 1.**
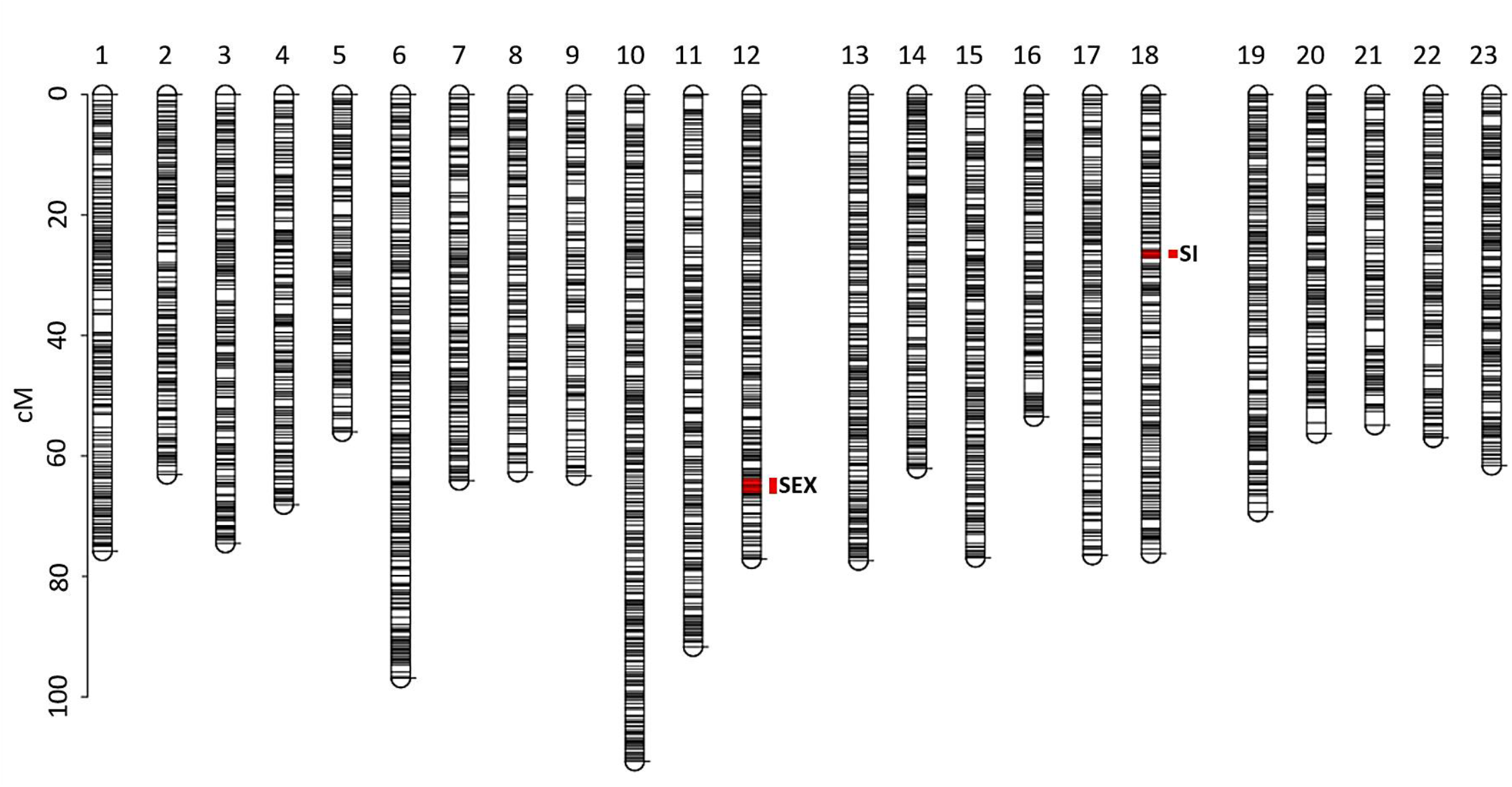
*Phillyrea angustifolia* sex-averaged linkage map showing the grouping and position of 15703 SNPs. The length of each of the 23 linkage groups is indicated by the vertical scale in cM. The markers strictly linked to sex and self-incompatibility (SI) phenotypes are shown in red. Markers that were clearly outliers at the end of some linkage groups were removed (see Table1, Figure S1).

For the comparison of male and hermaphrodites, an average of 44,565 SNPs were analyzed by SEX-DETector across the three subsamples, among which an average of 438 had a probability of following an XY pattern ≥0.80. We applied the same set of stringent filters and retained an average of 171 SNPs having been genotyped for at least 50% of the offspring, among which 41 had less than 5% of the offspring departing from the expected genotype under a XY model and were shared across the three subsets. Thirty-two of these SNPs followed a segregation pattern strictly consistent with a XY model. These 32 markers, corresponding to 8 loci, are distributed along a region of 2.216 cM on linkage group 12 (LG12, Figure 1) in the sex-averaged map. Relaxing the stringency or our thresholds, this region also contains five loci that strictly follow an XY segregation pattern but with fewer than 50% of offspring successfully genotyped, as well as 17 loci consistent with autosomal inheritance, possibly corresponding to polymorphisms accumulated within allelic lineages associated with either of the alternate sex phenotypes. Again, no SNP was found to follow a ZW pattern. This provides evidence that this independent region on LG12 is associated with sex, with a determination system akin to a XY system where males have the heterogametic genotype.

### Synteny analysis with the olive tree

About half (49%) of the 10,388 *P. angustifolia* loci used for the genetic map had a significant BLAST hit on the olive tree genome. Overall, the relative position of these hits was highly concordant with the structure of the linkage map. Indeed, the vast majority (79.7%) of loci belonging to a given linkage group had non-ambiguous matches on the same olive tree chromosome. Loci that did not follow this general pattern did not cluster on other chromosomes, suggesting either small rearrangements or mapping/assembly errors at the scale of individual loci. The order of loci within the linkage groups was also well conserved with only limited evidence for rearrangements (Figure 2, Figure 3), suggesting that the two genomes have remained largely collinear.

**Figure 2.**
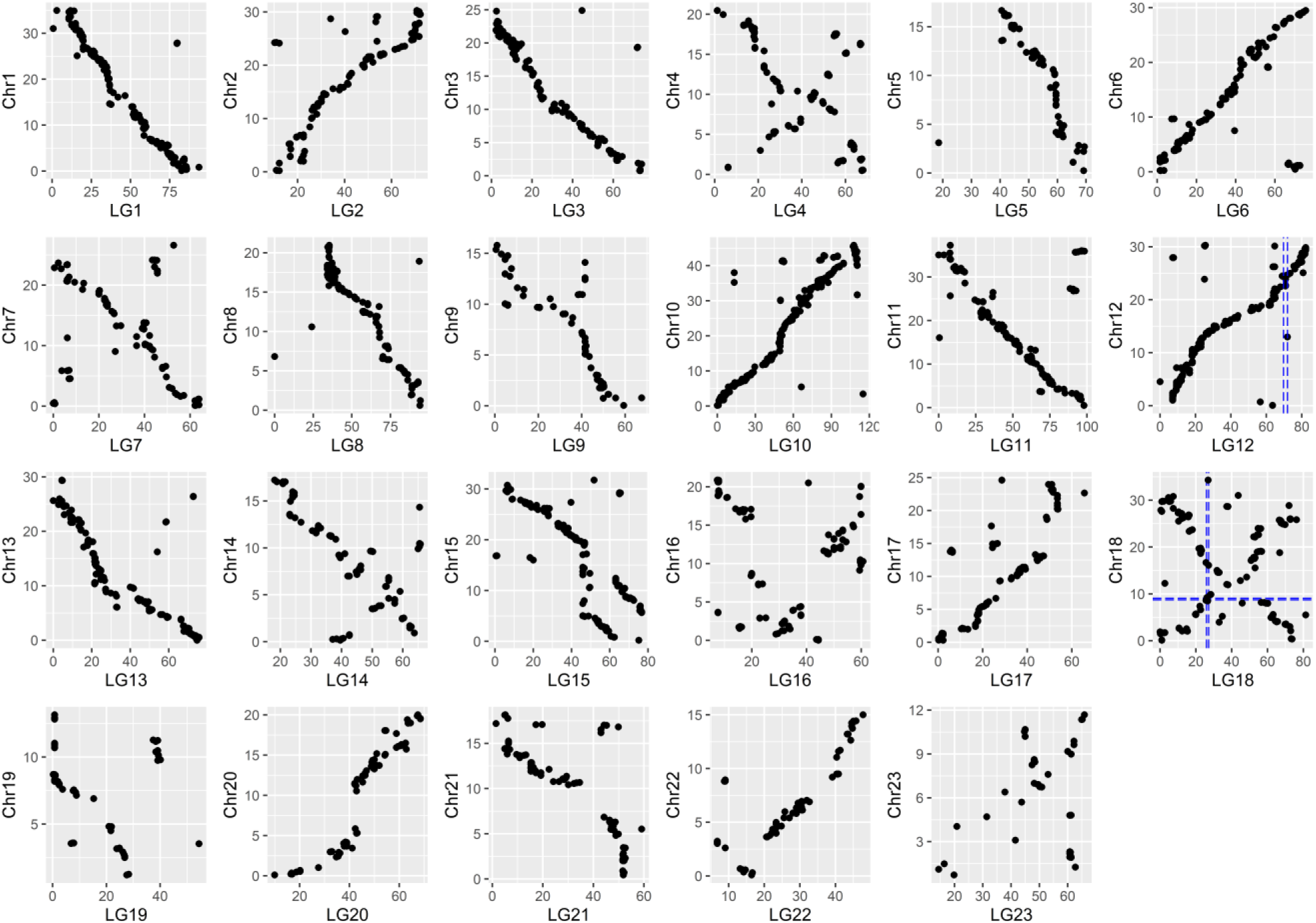
Synteny plot identifying homologous *P. angustifolia* linkage groups (LG, scale in cM) with olive tree chromosomes (Chr, scale in Mb). Lines connect markers in the *P. angustifolia* linkage map with their best BLAST hit in the *O. europea* genome and are colored according to the linkage group. Variation of the density of loci in bins of 3.125cM along linkage groups and 1 Mbp along chromosomes is shown in the inner circle as a black histogram.

**Figure 3.**
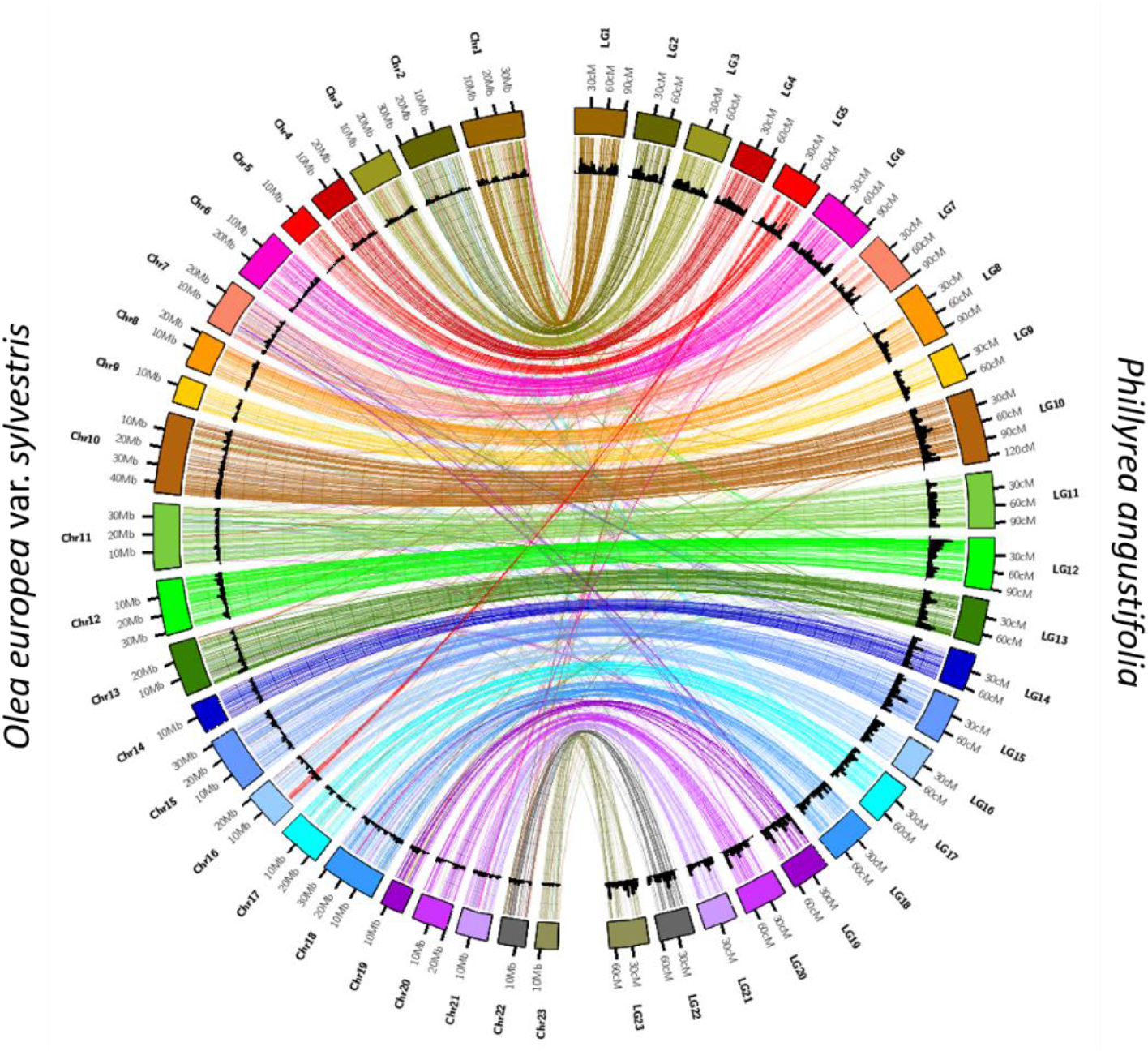
Visualization of chromosome-scale synteny by comparing the location of markers along the *P. angustifolia* linkage groups (LG, scale in cM) with the location of their best BLAST hit along the homologous olive tree chromosome (Chr, scale in Mbp). The vertical lines on LG12 and LG18 indicate the position of markers strictly associated with sex and SI phenotypes in *P. angustifolia*, respectively. The horizontal line on Chr18 indicates the chromosomal region containing the SI locus in *Olea europaea* according to Mariotti *et al.* (2020).

We then specifically inspected synteny between the linkage groups carrying either the sex or the SI locus and the olive tree genome (Figure 4). Synteny was good for LG12, the linkage group containing the markers associated with the sex phenotype. Among the 645 loci of LG12, 365 have good sequence similarity in the olive tree genome. Eighty eight percent had their best hits on the same chromosome of the olive tree (chromosome 12 per our numbering of the linkage groups), and the order of markers was largely conserved along this chromosome. Six loci contained in the region associated with sex on LG12 had hits on a single 1,940,009bp region on chromosome 12. This chromosomal interval contains 82 annotated genes in the olive tree genome (Table S1). In addition, eight loci in the sex region had their best hits on a series of five smaller scaffolds (Sca393, Sca1196, Sca1264, Sca32932, Sca969) that could not be reliably anchored in the main olive tree assembly but may nevertheless also contain candidates for sex determination. Collectively, these scaffolds represent 1.849.345bp of sequence in the olive tree genome and contain 57 annotated genes (Table S1).

**Figure 4.**
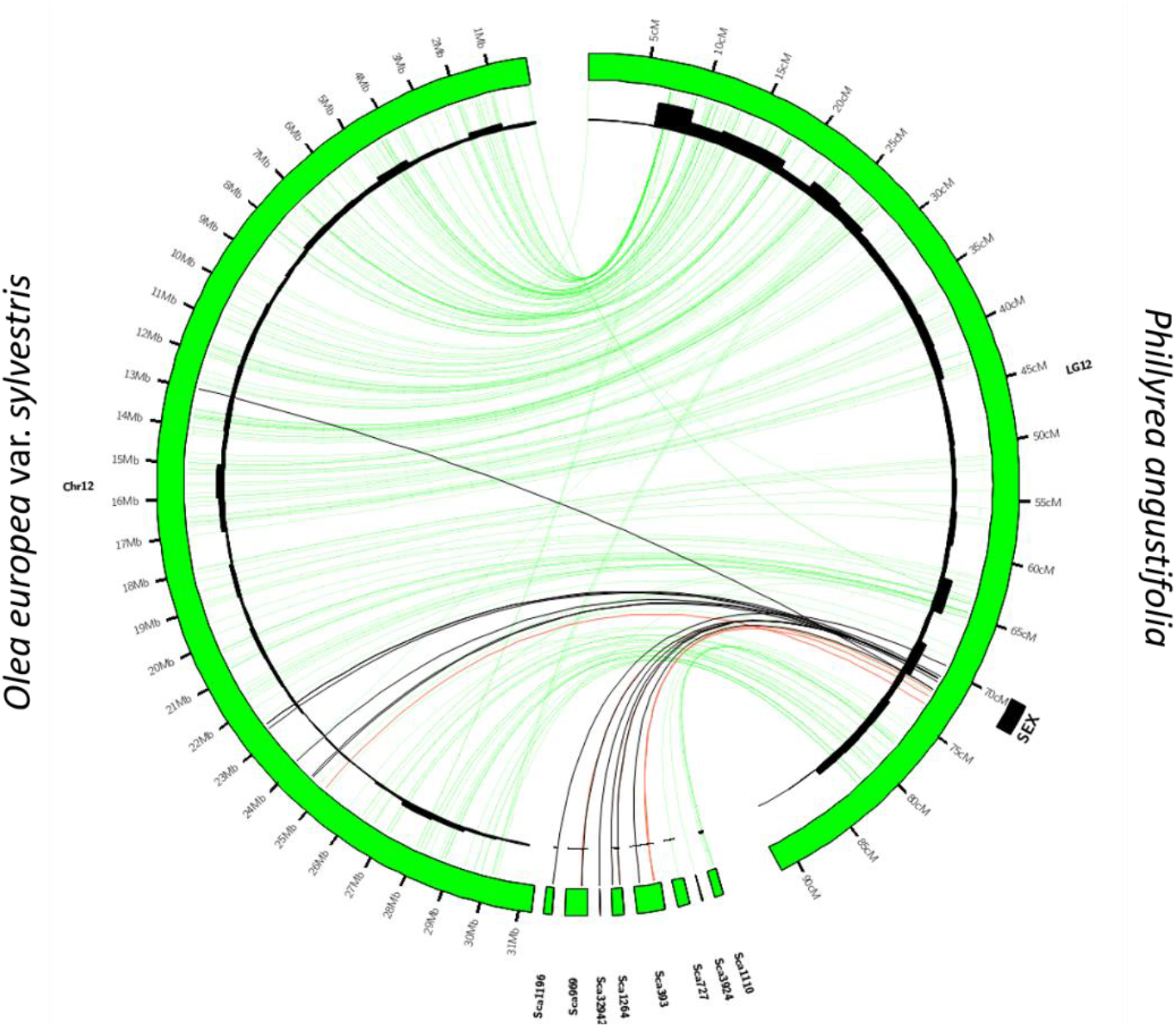
Synteny plot between the *P. angustifolia* linkage group 12 (scale in cM) and the olive tree chromosomes 12 and a series of unanchored scaffolds (scale in Mb). Lines connect markers in the *P. angustifolia* linkage map with their best BLAST hit in the *O. europea* genome. Green lines correspond to markers with autosomal inheritance. Black lines correspond to markers which strictly cosegregate with sex phenotypes (males *vs.* hermaphrodites). Red lines correspond to markers with strong but partial (95%) association with sex. Variation of the density of loci in bins of 3.125cM along linkage groups and 1 Mbp along chromosomes is shown in the inner circle as a black histogram.

Synteny was markedly poorer for markers on LG18, the linkage group containing the markers associated with the SI specificity phenotypes (Figure 5). Of the 440 loci on LG18, 203 had non-ambiguous BLAST hits on the olive tree genome. Although a large proportion (89%) had their best hits on chromosome 18, the order of hits along that chromosome suggested a large number of rearrangements. This more rearranged order was also observed for the six markers that were strictly associated with SI in *P. angustifolia*. Two of them had hits on a single region of 741,403bp on the olive tree genome. This region contains 32 annotated genes (Table S2) and contains two markers that were previously found to be genetically associated with SI directly in the olive tree by Mariotti *et al.* (2020). Three markers more loosely associated with SI in *P. angustifolia* had hits on a more distant region on chromosome 18 (19,284,909-19,758,630Mb). The three other strongly associated markers all had hits on scaffold 269, which contains 15 annotated genes and represents 545,128bp. Nine other loci strongly or loosely associated with SI had hits on a series of seven other unanchored scaffolds (Sca1199, Sca1200, Sca1287, Sca1579, Sca213, Sca327, Sca502) that collectively represent 96 annotated genes (Table S2) and 2,539,637bp.

**Figure 5.**
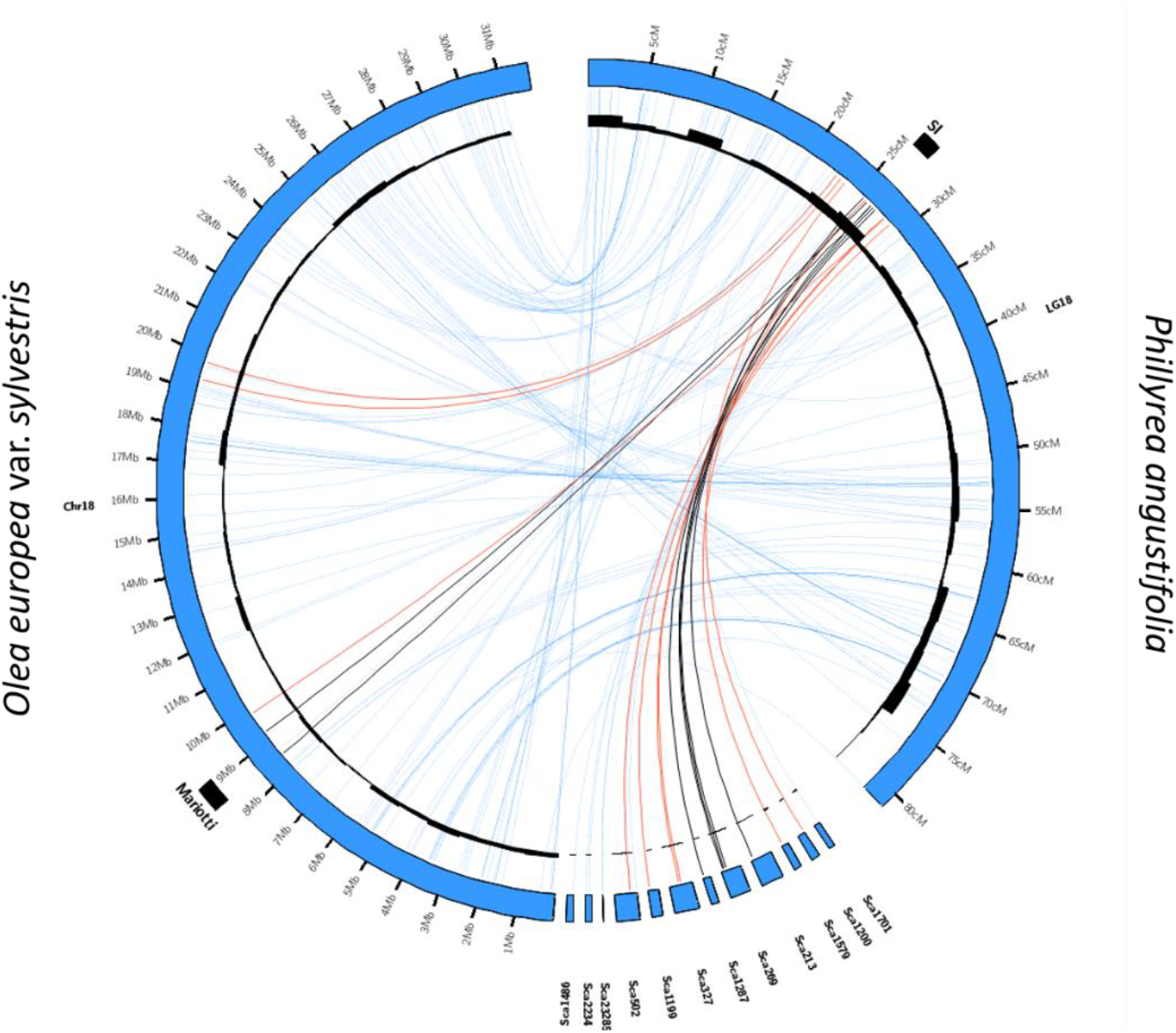
Synteny plot between the *P. angustifolia* linkage group 18 (scale in cM) and the olive tree chromosomes 18 and a series of unanchored scaffolds (scale in Mb). Lines connect markers in the *P. angustifolia* linkage map with their best BLAST hit in the *O. europea* genome. Blue lines correspond to markers with autosomal inheritance. Black lines correspond to markers which strictly cosegregate with SI phenotypes (Ha vs. Hb). Red lines correspond to markers with strong but partial (95%) association with SI. The region found to be genetically associated with SI in the olive tree by Mariotti et al. (2020) is shown by a black rectangle. Variation of the density of loci in bins of 3.125cM along linkage groups and 1 Mbp along chromosomes is shown in the inner circle as a black histogram.

## Discussion

Until now, studies have mostly relied on theoretical or limited genetic segregation analyses to investigate the evolution of sexual and SI phenotypes in *P. angustifolia* (Vassiliadis *et al.* 2002; Saumitou-Laprade *et al.* 2010; Husse *et al.* 2013; Billiard *et al.* 2015). In this study, we created the first genetic map of the androdioecious species *P. angustifolia* and identified the genomic regions associated with these two important reproductive phenotypes. The linkage map we obtained shows strong overall synteny with the olive tree genome, and reveals that sex and SI phenotypes segregate independently from one another, and are each strongly associated with a different genomic region (in LG18 and LG12, respectively).

The SI linked markers on LG18 are orthologous with the genomic interval recently identified by Mariotti *et al.* (2020) as the region controlling SI in the domesticated olive tree, providing strong reciprocal support that the determinants of SI are indeed located in this region. Interestingly, we observed a series of shorter scaffolds that could not previously be anchored in the main assembly of the olive tree genome but match genetic markers that are strictly linked to SI in *P. angustifolia.* These unanchored scaffolds provide a more complete set of genomic sequences that will be important to consider in the perspective of identifying the (currently elusive) molecular determinants of SI in these two species. We note that poor assembly of the S-locus region (Mariotti *et al.* 2020) was expected given the considerable levels of structural rearrangements typically observed in SI- and more generally in the mating type-determining regions (Goubet *et al.* 2012; Badouin *et al.* 2015), making *P. angustifolia* a useful resource to map the SI locus in the economically important species *O. europeae*.

Our observations also provide direct support to the hypothesis that the determinants of SI have remained at the same genomic position at least since the two lineages diverged, 30 to 40 Myrs ago (Besnard *et al.* 2009; Olofsson *et al.* 2019). Stability of the genomic location of SI genes has been observed in some Brassicaceae species, where the *SRK-SCR* system maps at orthologous positions in the Arabidopsis and Capsella genuses (Guo *et al.* 2011). In other Brassicaceae species, however, the SI system is found at different genomic locations, such as in Brassica and Leavenworthia. In the former, the molecular determinants have remained the same (also a series of *SRK-SCR* pairs, (Iwano *et al.* 2014), but in the latter SI seems to have evolved d*e novo* from exaptation of a pair of paralogous genes (Chantha *et al.* 2013; Chantha *et al.* 2017). Together with the fact that *P. angustifolia* pollen is able to trigger a robust SI response on *O. europaeae* stigmas (Saumitou-Laprade *et al.* 2017), our results provide strong support to the hypothesis that the *P. angustifolia* and *O. europaeae* SI systems are homologous. Whether mating type determinants occupy orthologous genomic regions in different species and rely on the same molecular players has also been discussed in oomycetes by Dussert *et al.* (2020).

Several approaches could now be used to refine the mapping of SI in *P. angustifolia,* and ultimately zero in on its molecular determinants. One possibility would require fine-mapping using larger offspring arrays, starting from our cross for which only a fraction of all phenotyped individuals were genotyped. Beyond the analysis of this controlled cross, evaluating whether the association of the SI phenotype still holds for markers within a larger set of accessions from diverse natural populations will constitute a powerful fine-mapping approach. Since the SI phenotypes seem to be functionally homologous across the Oleeae tribe (Vernet *et al.* 2016), the approach could, in principle, be extended to more distant SI species of the family like *L. vulgare* or *F. ornus*. Identification of sequences that have remained linked over these considerable time scales would represent excellent corroborative evidence to validate putative SI candidates. In parallel, an RNA-sequencing approach could be used to identify transcripts specific to the alternate SI phenotypes.

While comparison to the closely related *O. europeae* genome is a useful approach for the mapping of SI in *P. angustifolia*, it is *a priori* of limited use for mapping the sex-determining region, since the olive tree lineage has been entirely hermaphroditic for at least 32.22 Myrs (confidence interval: 28-36 Myrs) (FigS1 in Olofsson *et al.* 2019). Detailed exploration of the genomic region in the olive tree that is orthologous to the markers associated with sexual morphs in *P. angustifolia* is however interesting, as it may either have anciently played a role in sex determination and subsequently lost it, or alternatively it may contain quiescent sex-determining genes that have been activated specifically in *P. angustifolia*. At a broader scale, mapping and eventually characterizing the sex locus in other androdioecious species such as *F. ornus* could indicate whether the different instances of androdioecy in the family represent homologous phenotypes or independent evolutionary emergences.

Identifying the molecular mechanisms of the genes controlling SI and sex and tracing their evolution in a phylogenetic context would prove extremely useful. First, it could help understand the strong functional pleiotropy between sex and SI phenotypes, whereby males express universal SI compatibility (Saumitou-Laprade *et al.* 2010). In other words, males are able to transmit the SI specificities they inherited from their parents, but they do not express them themselves even though their pollen is fully functional. This intriguing feature of the SI system was key to solve the puzzle of why *P. angustifolia* maintains unusually high frequencies of males in natural populations (Husse *et al.* 2013), but the question of how being a male prevents expression of the SI phenotype in pollen is still open. A possibility is that the *M* allele of the sex locus contains a gene interacting negatively either with the pollen SI determinant itself or with a gene of the downstream response cascade. Identifying the molecular basis of this epistasis will be an interesting next step. Second, another intriguing feature of the system is segregation distortion, which is observed at several levels. Billiard *et al.* (2015) observed complete segregation bias in favor of males among the offspring of H_b_ hermaphrodites sired by males. Here, by phenotyping >1,000 offspring of a H_a_ hermaphrodite sired by a M_b_ male, we confirmed that this cross also entails a departure from Mendelian segregation, this time in favor of hermaphrodites, albeit of a lesser magnitude. Although the generality of this observation still remains to be determined by careful examination of the other possible crosses (H_a_ hermaphrodites × M_a_ and M_c_ males), it is clear that segregation distortion is a general feature of this system, as was already observed in other sex determination systems causing departures from equal sex ratios (e.g. Kozielska *et al.* 2010). Beyond identification of the mechanisms by which the distortions arise, pinpointing the evolutionary conditions leading to their emergence will be key to understanding the role they may have played in the evolution of this reproductive system.

More broadly, while sex and mating types are confounded in many species across the tree of life and cannot be distinguished, the question of when and how sex and mating types evolve separately raises several questions. The evolution of anisogamy (and hence, sexual differentiation) has been linked to that of mating types (Charlesworth 1978). In volvocine algae for instance, the mating-type locus in isogamous species is orthologous to the pair of U/V sex chromosomes in anisogamous/oogamous species, suggesting that the sex-determination system derives from the mating-type determination system (Geng *et al.* 2014). From this perspective the Oleaceae family is an interesting model system, where a SI system is ancestral, and in which some species have evolved sexual specialization that is aligned with the two SI phenotypes (e.g. in the polygamous *F. excelsior* males belong to the H_a_ SI group and can only mate with hermaphrodites or females of the H_b_ group, and the sexual system of *F. excelsior* can be viewed as subdioecy (Saumitou-Laprade *et al.* 2018). In other species, sexual phenotypes are disjoint from SI specificities and led to the differentiation of males and hermaphrodites. For instance, in the androdiecious *P. angustifolia* and probably *F. ornus*, the male determinant is genetically independent from the SI locus but fully linked to a genetic determinant causing the epistatic effect over SI (Billiard *et al.* 2015; Vernet *et al.* 2016). Yet other species have remained perfect hermaphrodites and have no trace of sexual differentiation whatsoever (*O. europeae*). Understanding why some species have followed one evolutionary trajectory while others have followed another will be an exciting avenue for future research (Billiard *et al.* 2011).

## Supporting information

Figure S1, linkage map with outliers SNPs. Figure S2, Comparison between the maternal and paternal genetic maps. Table S1 and S2, gene annotations

## Data accessibility

Fastq files for all 204 offspring and both parents are deposited in the [NCBI BioProject PRJNA724813](https://www.ncbi.nlm.nih.gov/bioproject/724813).

## Supplementary material

All scripts used can be accessed at https://github.com/Amelie-Carre/Genetic-map-of-Phillyrea-angustifolia.

## Acknowledgements

We thank Sylvain Bertrand and Fantin Carpentier for technical help for the phenotyping and Jos Käfer for scientific discussion and help in applying Sex-DETector to our material. We thank Jacques Lepart†, Mathilde Dufay, Pierre Olivier Cheptou, Xavier Vekemans, Sylvain Billiard and Bénédicte Felter for scientific discussions. Field and laboratory work for phenotyping were done at the Platform Terrains d’Expériences (Labex CeMEB ANR-10-LABX-0004-CeMEB) of the Centre d’Ecologie Fonctionnelle et Evolutive (CEFE, CNRS) with the help of Thierry Mathieu and David Degueldre and at the platform Serres cultures et terrains expérimentaux of the Lille University with the help of Nathalie Faure and Angélique Bourceaux. We are grateful to Marie-Pierre Dubois for providing access to the microscopy facilities at the SMGE (Service des Marqueurs Génétiques en Ecologie) platform (CEFE). This work was funded by the French National Research Agency through the project ‘TRANS’ (ANR-11-BSV7-013-03) and by a grant from the European Research Council (NOVEL project, grant #648321). A.C was supported by a doctoral grant from the French ministry of research. The authors also thank the Région Hauts-de-France, and the Ministère de l’Enseignement Supérieur et de la Recherche (CPER Climibio), and the European Fund for Regional Economic Development for their financial support. We also thank the HPC Computing Mésocentre of the University of Lille which provided us with the computing grid. We thank Tatiana Giraud, Ricardo C. Rodríguez de la Vega and two anonymous referees for constructive reviews of an earlier version. Version 7 of this preprint has been peer-reviewed and recommended by Peer Community In Genomics (https://doi.org/10.24072/pci.genomics.100011).

## Conflict of interest disclosure and Author Contributions

The authors of this preprint declare that they have no financial conflict of interest with the content of this article. All authors contributed to the study presented in this paper. PS-L and PV developed, designed and oversaw the study; they coordinated the cross and carried out the phenotyping and stigma tests. CG performed the seedling paternity analysis. SS performed DNA extraction, library preparation and organized sequencing. AC and SG constructed the data analysis pipeline and AC, SS, PS-L and VC interpreted the results and wrote the manuscript.

## Appendix

File name: supplementary_file

File format: Portable document format (pdf)

Title of data:

Figure S1. *Phillyrea angustifolia* sex-averaged linkage map showing the grouping and position of 15812 SNPs.

Figure S2. Comparison between the maternal and paternal genetic maps.

Table S1. List of the 82 gene annotations in the chromosomal interval of the olive tree genome bounded by *P. angustifolia* loci strictly associated with sex.

Table S2. List of the 32 gene annotations in the chromosomal interval of the olive tree genome bounded by *P. angustifolia* loci strictly associated with SI.

## Notes

https://github.com/Amelie-Carre/Genetic-map-of-Phillyrea-angustifolia

http://www.ncbi.nlm.nih.gov/bioproject/724813

### Competing Interest Statement

The authors have declared no competing interest.

### Summary of Updates

Version 7 of this preprint has been peer-reviewed and recommended by Peer Community In Genomics (https://doi.org/10.24072/pci.genomics.100011)

## References

Badouin, H., M. E. Hood, J. Gouzy, G. Aguileta, S. Siguenza et al., 2015 Chaos of rearrangements in the mating-type chromosomes of the anther-smut fungus *Microbotryum lychnidis-dioicae*. Genetics 200: 1275–1284. https://dx.doi.org/10.1534/genetics.115.177709

Barrett, S. C., 2002 The evolution of plant sexual diversity. Nature Reviews Genetics 3: 274–284. https://dx.doi.org/10.1038/nrg776

Barrett, S. C. H., 1990 The evolution and adaptive significance of heterostyly. Trends in Ecology & Evolution 5: 144–148. https://doi.org/10.1016/0169-5347(90)90220-8

Barrett, S. C. H., 1992 Heterostylous genetic polymorphisms: Model systems for evolutionary analysis, pp. 1–29 in Evolution and Function of Heterostyly, edited by S. C. H. Barrett. Springer Berlin Heidelberg, Berlin, Heidelberg. https://dx.doi.org/10.1007/978-3-642-86656-2_1

Barrett, S. C. H., 1998 The evolution of mating strategies in flowering plants. Trends in Plant Science 3: 335—341. http://dx.doi.org/10.1016/S1360-1385(98)01299-0

Barrett, S. C. H., 2019 ‘A most complex marriage arrangement’: recent advances on heterostyly and unresolved questions. New Phytologist 224: 1051–1067. https://doi.org/10.1111/nph.16026

Besnard, G., R. Rubio de Casas, P.-A. Christin and P. Vargas, 2009 Phylogenetics of *Olea* (Oleaceae) based on plastid and nuclear ribosomal DNA sequences: Tertiary climatic shifts and lineage differentiation times. Annals of Botany 104: 143–160. https://dx.doi.org/10.1093/aob/mcp105

Billiard, S., L. Husse, P. Lepercq, C. Gode, A. Bourceaux et al., 2015 Selfish male-determining element favors the transition from hermaphroditism to androdioecy. Evolution 69: 683–693. https://dx.doi.org/10.1111/evo.12613

Billiard, S., M. López-Villavicencio, B. Devier, M. E. Hood, C. Fairhead et al., 2011 Having sex, yes, but with whom? Inferences from fungi on the evolution of anisogamy and mating types. Biological Reviews 86: 421–442. https://doi.org/10.1111/j.1469-185X.2010.00153.x

Castric, V., and X. Vekemans, 2004 Plant self-incompatibility in natural populations: a critical assesment of recent theoretical and empirical advances. Molecular Ecology 13: 2873–2889. https://dx.doi.org/10.1111/j.1365-294X.2004.02267.x

Catchen, J. M., A. Amores, P. Hohenlohe, W. Cresko and J. H. Postlethwait, 2011 Stacks: building and genotyping loci de novo from short-read sequences. G3 (Bethesda, Md.) 1: 171–182. https://dx.doi.org/10.1534/g3.111.000240

Chantha, S.-C., A. C. Herman, V. Castric, X. Vekemans, W. Marande et al., 2017 The unusual *S* locus of *Leavenworthia* is composed of two sets of paralogous loci. New Phytologist 216: 1247–1255. https://doi.org/10.1111/nph.14764

Chantha, S.-C., A. C. Herman, A. E. Platts, X. Vekemans and D. J. Schoen, 2013 Secondary evolution of a self-incompatibility locus in the Brassicaceae *Genus leavenworthia*. PLOS Biology 11: e1001560. https://dx.doi.org/10.1371/journal.pbio.1001560

Charlesworth, B., 1978 The population genetics of anisogamy. Journal of Theoretical Biology 73: 347–357. https://doi.org/10.1016/0022-5193(78)90195-9

Charlesworth, D., and B. Charlesworth, 1979 A model for the evolution of distyly. The American Naturalist 114: 467–498. https://doi.org/10.1086/283496

De-Kayne, R., and P. G. D. Feulner, 2018 A european whitefish linkage map and its implications for understanding genome-wide synteny between salmonids following whole genome duplication. G3 (Bethesda) 8: 3745–3755. https://dx.doi.org/10.1534/g3.118.200552

De Cauwer, I., P. Vernet, S. Billiard, C. Godé, A. Bourceaux et al., 2020 Widespread coexistence of self-compatible and self-incompatible phenotypes in a diallelic self-incompatibility system in *Ligustrum vulgare* (Oleaceae). bioRxiv : 2020.2003.2026.009399. https://dx.doi.org/10.1101/2020.03.26.009399

De Nettancourt, D., 1977 Incompatibility in Angiosperms. Springer-Verlag, Berlin, Heidelberg et New York.

Diggle, P. K., V. S. Di Stilio, A. R. Gschwend, E. M. Golenberg, R. C. Moore et al., 2011 Multiple developmental processes underlie sex differentiation in angiosperms. Trends in Genetics 27: 368–376. https://doi.org/10.1016/j.tig.2011.05.003

Dommée, B., J. D. Thompson and F. Cristini, 1992 Distylie chez *Jasminum fruticans* L.: hypothèse de la pollinisation optimale basée sur les variations de l’écologie intraflorale. Bulletin de la Société Botanique de France. Lettres Botaniques 139: 223–234. https://dx.doi.org/10.1080/01811797.1992.10824960

Dulberger, R., 1992 Floral polymorphisms and their functional significance in the heterostylous syndrome, pp. 41–84 in Evolution and Function of Heterostyly, edited by S. C. H. Barrett. Springer Berlin Heidelberg, Berlin, Heidelberg. https://dx.doi.org/10.1007/978-3-642-86656-2_3

Dupin, J., P. Raimondeau, C. Hong-Wa, S. Manzi, M. Gaudeul et al., 2020 Resolving the phylogeny of the olive family (Oleaceae): Confronting Information from organellar and nuclear genomes. Genes 11: 1508. https://dx.doi.org/10.3390/genes11121508

Dussert, Y., L. Legrand, I. D. Mazet, C. Couture, M.-C. Piron et al., 2020 Identification of the first oomycete mating-type locus sequence in the grapevine downy mildew pathogen, *Plasmopara viticola*. Current biology : CB 30: 3897–3907.e3894. https://dx.doi.org/10.1016/j.cub.2020.07.057

Geng, S., P. De Hoff and J. G. Umen, 2014 Evolution of sexes from an ancestral mating-type specification pathway. PLoS biology 12: e1001904–e1001904. https://dx.doi.org/10.1371/journal.pbio.1001904

Goubet, P. M., H. Bergès, A. Bellec, E. Prat, N. Helmstetter et al., 2012 Contrasted patterns of molecular evolution in dominant and recessive self-incompatibility haplotypes in *Arabidopsis*. PLOS Genetics 8: e1002495. https://dx.doi.org/10.1371/journal.pgen.1002495

Guo, Y.-L., X. Zhao, C. Lanz and D. Weigel, 2011 Evolution of the S-locus region in *Arabidopsis* relatives. Plant physiology 157: 937–946. https://dx.doi.org/10.1104/pp.111.174912

Husse, L., S. Billiard, J. Lepart, P. Vernet and P. Saumitou-Laprade, 2013 A one-locus model of androdioecy with two homomorphic self-incompatibility groups: expected *vs.* observed male frequencies. Journal of evolutionary biology 26: 1269–1280. https://dx.doi.org/10.1111/jeb.12124

Iwano, M., K. Ito, H. Shimosato-Asano, K.-S. Lai and S. Takayama, 2014 Self-incompatibility in the Brassicaceae, pp. 245–254 in Sexual Reproduction in Animals and Plants, edited by H. Sawada, N. Inoue and M. Iwano. Springer Japan, Tokyo. https://doi.org/10.1007/978-4-431-54589-7_21

Iwano, M., and S. Takayama, 2012 Self/non-self discriminition in angiosperm self-incompatibility. Current Opinion in Plant Biology 15: 78–83. https://doi.org/10.1016/j.pbi.2011.09.003

Jiménez-Ruiz, J., J. A. Ramírez-Tejero, N. Fernández-Pozo, M. d. l. O. Leyva-Pérez, H. Yan et al., 2020 Transposon activation is a major driver in the genome evolution of cultivated olive trees (*Olea europaea L.*). The Plant Genome 13: e20010. https://doi.org/10.1002/tpg2.20010

Keller, B., J. D. Thomson and E. Conti, 2014 Heterostyly promotes disassortative pollination and reduces sexual interference in Darwin’s primroses: evidence from experimental studies. Functional Ecology 28: 1413–1425. https://doi.org/10.1111/1365-2435.12274

Kozielska, M., F. J. Weissing, L. W. Beukeboom and I. Pen, 2010 Segregation distortion and the evolution of sex-determining mechanisms. Heredity 104: 100–112. https://dx.doi.org/10.1038/hdy.2009.104

Krzywinski, M. I., J. E. Schein, I. Birol, J. Connors, R. Gascoyne et al., 2009 Circos: An information aesthetic for comparative genomics. Genome Research. https://dx.doi.org/10.1101/gr.092759.109

Kubo, K.-i., T. Paape, M. Hatakeyama, T. Entani, A. Takara et al., 2015 Gene duplication and genetic exchange drive the evolution of S-RNase-based self-incompatibility in *Petunia*. Nature Plants 1: 1–9. https://doi.org/10.1038/nplants.2014.5

Langmead, B., and S. L. Salzberg, 2012 Fast gapped-read alignment with Bowtie 2. Nature methods 9: 357–359. https://dx.doi.org/10.1038/nmeth.1923

Li, H., and R. Durbin, 2009 Fast and accurate short read alignment with Burrows-Wheeler transform. Bioinformatics 25: 1754–1760. https://dx.doi.org/10.1093/bioinformatics/btp324

Li, J., J. M. Cocker, J. Wright, M. A. Webster, M. McMullan et al., 2016 Genetic architecture and evolution of the S locus supergene in *Primula vulgaris*. Nature plants 2: 16188. https://dx.doi.org/10.1038/nplants.2016.188

Lundqvist, A., 1956 Self-incompatibility in rye. Hereditas 42: 293–348. https://doi.org/10.1111/j.1601-5223.1956.tb03021.x

Mariotti, R., S. Pandolfi, I. De Cauwer, P. Saumitou-Laprade, P. Vernet et al., 2020 Diallelic self-incompatibility is the main determinant of fertilization patterns in olive orchards. Evolutionary Applications n/a. https://doi.org/10.1111/eva.13175

Muyle, A., J. Kafer, N. Zemp, S. Mousset, F. Picard et al., 2016 SEX-DETector: A probabilistic approach to study sex chromosomes in non-model organisms. Genome Biol Evol 8: 2530–2543. https://dx.doi.org/10.1093/gbe/evw172

Olofsson, J. K., I. Cantera, C. Van de Paer, C. Hong-Wa, L. Zedane et al., 2019 Phylogenomics using low-depth whole genome sequencing: A case study with the olive tribe. Molecular Ecology Resources 19: 877–892. https://doi.org/10.1111/1755-0998.13016

Pannell, J. R., and G. Korbecka, 2010 Mating-system evolution: Rise of the irresistible males. Current Biology 20: R482–R484. https://doi.org/10.1016/j.cub.2010.04.033

Pannell, J. R., and M. Voillemot, 2015 Plant mating systems: female sterility in the driver’s seat. Current Biology 25: R511–514. https://dx.doi.org/10.1016/j.cub.2015.04.044

Peterson, B. K., J. N. Weber, E. H. Kay, H. S. Fisher and H. E. Hoekstra, 2012 Double digest RADseq: an inexpensive method for de novo SNP discovery and genotyping in model and non-model species. PLoS One 7: e37135. https://dx.doi.org/10.1371/journal.pone.0037135

Rastas, P., 2017 Lep-MAP3: robust linkage mapping even for low-coverage whole genome sequencing data. Bioinformatics 33: 3726–3732. https://dx.doi.org/10.1093/bioinformatics/btx494

Rochette, N. C., and J. M. Catchen, 2017 Deriving genotypes from RAD-seq short-read data using Stacks. Nature Protocols 12: 2640–2659. https://dx.doi.org/10.1038/nprot.2017.123

Sakai, A. K., and S. G. Weller, 1999 Gender and sexual dimorphism in flowering plants: A review of terminology, biogeographic patterns, ecological correlates, and phylogenetic approaches, pp. 1–31 in Gender and Sexual Dimorphism in Flowering Plants, edited by M. A. Geber, T. E. Dawson and L. F. Delph. Springer Berlin Heidelberg, Berlin, Heidelberg. https://dx.doi.org/10.1007/978-3-662-03908-3_1

Saumitou-Laprade, P., P. Vernet, A. Dowkiw, S. Bertrand, S. Billiard et al., 2018 Polygamy or subdioecy? The impact of diallelic self-incompatibility on the sexual system in *Fraxinus excelsior* (Oleaceae). Proceedings. Biological sciences 285: 20180004. https://dx.doi.org/10.1098/rspb.2018.0004

Saumitou-Laprade, P., P. Vernet, C. Vassiliadis, Y. Hoareau, G. de Magny et al., 2010 A self-incompatibility system explains high male frequencies in an androdioecious plant. Science 327: 1648–1650. https://dx.doi.org/10.1126/science.1186687

Saumitou-Laprade, P., P. Vernet, X. Vekemans, S. Billiard, S. Gallina et al., 2017 Elucidation of the genetic architecture of self-incompatibility in olive: Evolutionary consequences and perspectives for orchard management. Evolutionary Applications 1–14. https://dx.doi.org/10.1111/eva.12457

Taylor, H., 1945 Cyto-taxonomy and phylogeny of the oleaceae. Brittonia 5: 337–367. https://dx.doi.org/10.2307/2804889

Tsagkogeorga, G., V. Cahais and N. Galtier, 2012 The population genomics of a fast evolver: high levels of diversity, functional constraint, and molecular adaptation in the tunicate *Ciona intestinalis*. Genome Biol Evol 4: 740–749. https://dx.doi.org/10.1093/gbe/evs054

Unver, T., Z. Wu, L. Sterck, M. Turktas, R. Lohaus et al., 2017 Genome of wild olive and the evolution of oil biosynthesis. Proceedings of the National Academy of Sciences of the United States of America 114: E9413–e9422. https://dx.doi.org/10.1073/pnas.1708621114

Vassiliadis, C., P. Saumitou-Laprade, J. Lepart and F. Viard, 2002 High male reproductive success of hermaphrodites in the androdioecious *Phillyrea angustifolia*. Evolution 56: 1362–1373. https://doi.org/10.1111/j.0014-3820.2002.tb01450.x

Vernet, P., P. Lepercq, S. Billiard, A. Bourceaux, J. Lepart et al., 2016 Evidence for the long-term maintenance of a rare self-incompatibility system in Oleaceae. New Phytologist 210: 1408–1417. https://dx.doi.org/10.1111/nph.13872

Wallander, E., and V. A. Albert, 2000 Phylogeny and classification of Oleaceae based on rps16 and trnL-F sequence data. American Journal of Botany 87: 1827–1841. https://doi.org/10.2307/2656836

Wright, S., 1939 The distribution of self-sterility alleles in populations. Genetics 24: 538–552.

