## Supplementary material for "Genetic mapping of sex and self-incompatibility determinants in the androdioecious plant *Phillyrea angustifolia*": Figure S1, linkage map with outliers SNPs. Figure S2, Comparison between the maternal and paternal genetic maps. Table S1 and S2, gene annotations

Figure S1. *Phillyrea angustifolia* sex-averaged linkage map showing the grouping and position of 15812 SNPs. The length of each of the 23 linkage groups is indicated by the vertical scale in cM. The markers strictly linked to sex and self-incompatibility (SI) phenotypes are shown in red.

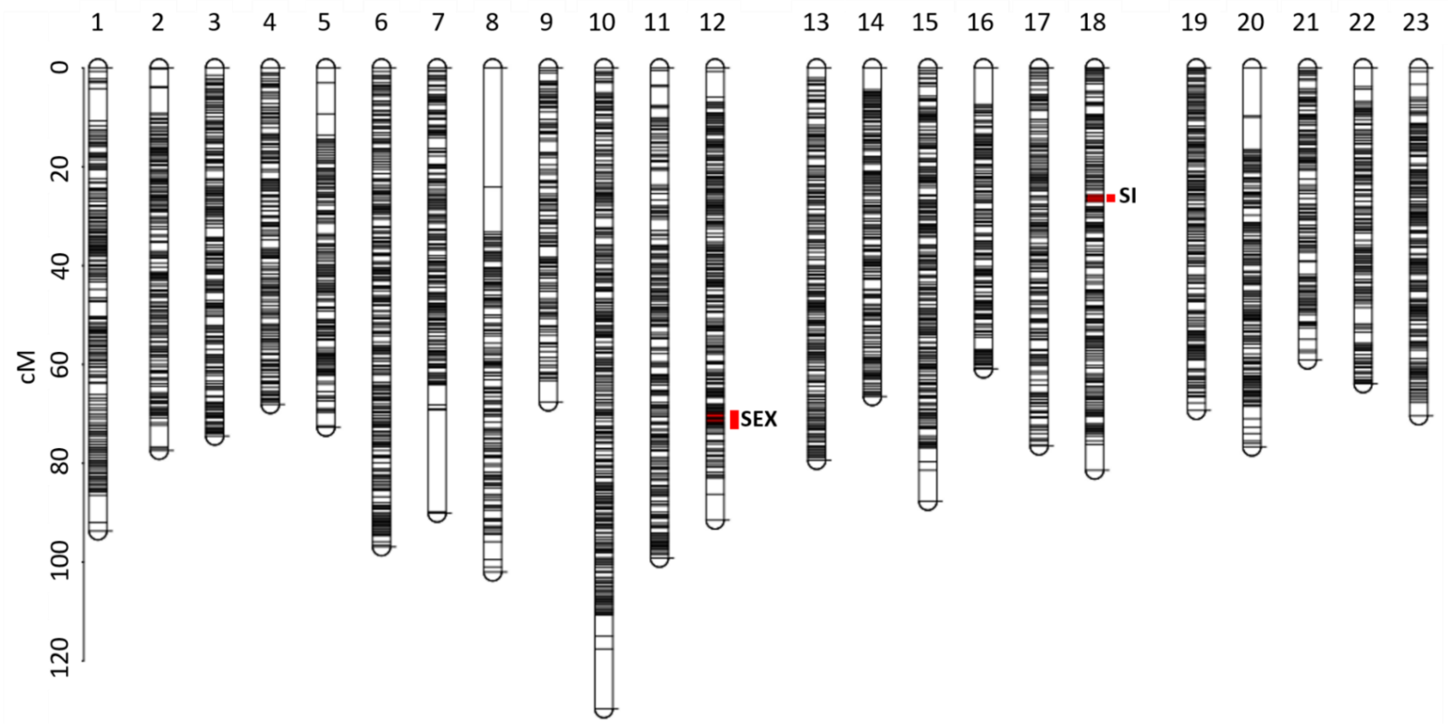

Figure S2. Comparison between the maternal and paternal genetic maps. The vertical and horizontal lines on LG12 and LG18 indicate the position of markers strictly associated with sex and SI phenotypes.

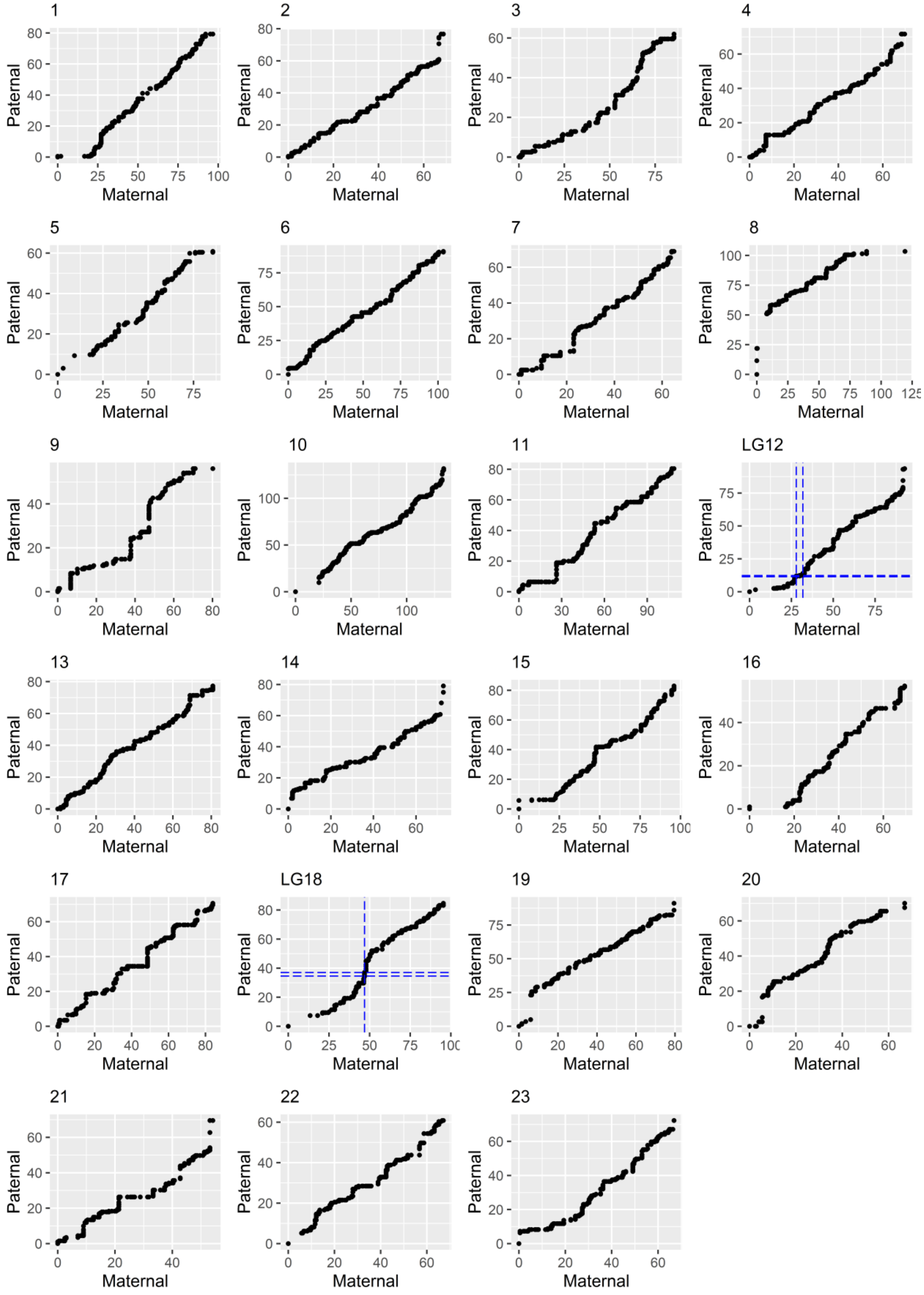

Table S1. List of the 82 gene annotations in the chromosomal interval of the olive tree genome bounded by *P. angustifolia* loci strictly associated with sex. This list is completed by the 57 gene annotations in the five scaffolds containing loci strongly or loosely associated with sex in *P. angustifolia*. The description field is taken from Unver *et al.* (2020) and is based on automated blast annotation. NA corresponds to predicted gene models with no hit.

| genome location | Sequence ID | Description | GO annotation |
|---|---|---|---|
| chr12 | Oeu003179.1 | ribonuclease h | |
| chr12 | Oeu003182.1 | 50S ribosomal protein L15, chloroplastic | F:GO:0003735:structural constituent of ribosome ; P:GO:0006412:translation; C:GO:0015934:large ribosomal subunit |
| chr12 | Oeu003184.1 | caffeoylshikimate esterase | |
| chr12 | Oeu003185.3 | uncharacterized protein LOC111406129 | |
| chr12 | Oeu003188.1 | uncharacterized protein LOC111406130 | C:GO:0016020:membrane ; F:GO:0016757:glycosyltransferase activity |
| chr12 | Oeu003330.1 | uncharacterized protein LOC111406115 isoform X1 | P:GO:0015031:protein transport |
| chr12 | Oeu003331.1 | probable NADH dehydrogenase [ubiquinone] 1 alpha subcomplex subunit 5, mitochondrial | P:GO:0022904:respiratory electron transport chain |
| chr12 | Oeu003332.1 | late embryogenesis abundant protein At1g64065-like | |
| chr12 | Oeu003334.1 | reticulon-like protein B5 | |
| chr12 | Oeu003335.2 | WD repeat-containing protein 91 homolog | F:GO:0005515:protein binding |
| chr12 | Oeu003336.1 | zinc finger MYM-type protein 1-like | |
| chr12 | Oeu003337.1 | probable WRKY transcription factor 19 | |
| chr12 | Oeu003338.1 | putative SNAP25 homologous protein SNAP30 | |
| chr12 | Oeu003339.1 | Hypothetical predicted protein | |
| chr12 | Oeu003341.1 | uncharacterized protein LOC111406154 isoform X2 | |
| chr12 | Oeu003342.1 | ABC transporter C family member 10-like | F:GO:0005524:ATP binding; C:GO:0016021:integral component of membrane; F:GO:0042626:ATPase-coupled transmembrane transporter activity; P:GO:0055085:transmembrane transport |
| chr12 | Oeu003343.1 | uncharacterized protein LOC111394170 | P:GO:0000723:telomere maintenance; F:GO:0003678:DNA helicase activity; P:GO:0006281:DNA repair |
| chr12 | Oeu003344.1 | probable leucine-rich repeat receptor kinase At1g68400 | F:GO:0004672:protein kinase activity; F:GO:0005524:ATP binding; P:GO:0006468:protein phosphorylation |
| chr12 | Oeu003345.3 | probable LRR receptor-like serine/threonine-protein kinase At1g51880 | F:GO:0004672:protein kinase activity; F:GO:0005524:ATP binding; P:GO:0006468:protein phosphorylation; F:GO:0046872:metal ion binding |
| chr12 | Oeu003346.1 | probable inactive receptor kinase At4g23740 | F:GO:0004672:protein kinase activity; F:GO:0005524:ATP binding; P:GO:0006468:protein phosphorylation |
| chr12 | Oeu003347.1 | pollen receptor-like kinase 1 | F:GO:0004672:protein kinase activity; F:GO:0005524:ATP binding; P:GO:0006468:protein phosphorylation |

| | | | |
|---|---|---|---|
| chr12 | Oeu003348.1 | probable inactive receptor kinase At4g23740 | F:GO:0004672:protein kinase activity; F:GO:0005524:ATP binding; P:GO:0006468:protein phosphorylation |
| chr12 | Oeu010693.1 | phytolongin Phyl2.1-like | C:GO:0016021:integral component of membrane |
| chr12 | Oeu010694.1 | protein RKD3 | F:GO:0003700:DNA-binding transcription factor activity |
| chr12 | Oeu010695.1 | nephrocystin-3 | F:GO:0005515:protein binding |
| chr12 | Oeu010696.3 | hypothetical protein H5410_017114 | P:GO:0006606:protein import into nucleus; F:GO:0031267:small GTPase binding |
| chr12 | Oeu010697.1 | uncharacterized protein LOC111406111 | |
| chr12 | Oeu010698.1 | Hypothetical predicted protein | |
| chr12 | Oeu010700.1 | thylakoid lumenal 17.4 kDa protein, chloroplastic-like isoform X1 | |
| chr12 | Oeu010701.1 | Hypothetical predicted protein | |
| chr12 | Oeu010702.1 | WD repeat-containing protein YMR102C-like isoform X1 | F:GO:0005515:protein binding |
| chr12 | Oeu010703.1 | pentatricopeptide repeat-containing protein At2g13600-like | F:GO:0005515:protein binding |
| chr12 | Oeu010704.1 | protein yippee-like At4g27745 | |
| chr12 | Oeu010705.1 | oligopeptide transporter 6-like | P:GO:0055085:transmembrane transport |
| chr12 | Oeu010706.1 | molybdate-anion transporter-like | F:GO:0015098:molybdate ion transmembrane transporter activity; P:GO:0015689:molybdate ion transport; C:GO:0016021:integral component of membrane |
| chr12 | Oeu010708.1 | receptor-like protein 12 | F:GO:0005515:protein binding |
| chr12 | Oeu010709.1 | pentatricopeptide repeat-containing protein At3g18020 | F:GO:0005515:protein binding |
| chr12 | Oeu010710.1 | protein PHOSPHATE STARVATION RESPONSE 3-like | |
| chr12 | Oeu010711.1 | protein PHOSPHATE STARVATION RESPONSE 1-like | |
| chr12 | Oeu010714.1 | uncharacterized protein LOC111377053 | P:GO:0006508:proteolysis; F:GO:0008234:cysteine-type peptidase activity |
| chr12 | Oeu010715.1 | Hypothetical predicted protein | |
| chr12 | Oeu010716.1 | uncharacterized protein LOC111407041 | |
| chr12 | Oeu010717.1 | uncharacterized protein LOC111395991 | |
| chr12 | Oeu015503.1 | nudix hydrolase 25-like | F:GO:0016787:hydrolase activity |
| chr12 | Oeu015505.1 | uncharacterized protein LOC111406139 isoform X2 | |
| chr12 | Oeu015506.1 | uncharacterized protein LOC111406809 | |
| chr12 | Oeu015507.1 | protein C2-DOMAIN ABA-RELATED 7-like isoform X1 | |

| | | | |
|---|---|---|---|
| chr12 | Oeu015509.1 | uncharacterized protein LOC111390401 | |
| chr12 | Oeu015510.1 | probable CCR4-associated factor 1 homolog 11 | F:GO:0003676; F:GO:0004535; C:GO:0030014 |
| chr12 | Oeu015511.1 | uncharacterized protein LOC111406141 | |
| chr12 | Oeu015512.1 | inositol-tetrakisphosphate 1-kinase 1-like | F:GO:0000287:magnesium ion binding; F:GO:0005524:ATP binding; P:GO:0032957:inositol trisphosphate metabolic process; F:GO:0047325:inositol tetrakisphosphate 1-kinase activity; F:GO:0052725:inositol-1,3,4-trisphosphate 6-kinase; F:GO:0052726:inositol-1,3,4-trisphosphate 5-kinase |
| chr12 | Oeu032517.1 | 5 -3 exoribonuclease 3 | |
| chr12 | Oeu032519.1 | uncharacterized protein LOC111406806 | F:GO:0003677:DNA binding; F:GO:0046983:protein dimerization activity |
| chr12 | Oeu032520.1 | Hypothetical predicted protein | |
| chr12 | Oeu032521.1 | uncharacterized protein LOC111406134 | F:GO:0003677:DNA binding; F:GO:0046983:protein dimerization activity |
| chr12 | Oeu032522.1 | ATP-dependent DNA helicase 2 subunit KU80-like | |
| chr12 | Oeu032523.1 | uncharacterized protein LOC111373134 | |
| chr12 | Oeu037452.1 | ---NA--- | |
| chr12 | Oeu037453.1 | ---NA--- | |
| chr12 | Oeu037454.1 | nucleolar complex protein 2 homolog | |
| chr12 | Oeu037965.1 | uncharacterized protein LOC111406151 isoform X2 | |
| chr12 | Oeu037967.1 | GRF1-interacting factor 2-like | F:GO:0003713:transcription coactivator activity |
| chr12 | Oeu037968.1 | Hypothetical predicted protein | |
| chr12 | Oeu037969.1 | uncharacterized protein LOC111406816 | C:GO:0000127:transcription factor TFIIIC complex; F:GO:0004402:histone acetyltransferase activity; F:GO:0005515:protein binding; P:GO:0006384:transcription initiation from RNA polymerase III promoter |
| chr12 | Oeu037970.1 | uncharacterized protein LOC111392799 | |
| chr12 | Oeu037972.1 | uncharacterized protein LOC111404605 | |
| chr12 | Oeu037973.2 | polyadenylate-binding protein-interacting protein 12-like | F:GO:0003676:nucleic acid binding |
| chr12 | Oeu037975.2 | uncharacterized protein LOC111406147 | |
| chr12 | Oeu037977.2 | uncharacterized protein LOC111406815 | P:GO:0006629:lipid metabolic process; F:GO:0008970:phospholipase A1 activity |
| chr12 | Oeu037979.1 | nascent polypeptide-associated complex subunit alpha-like protein 2 | C:GO:0005854:nascent polypeptide-associated complex |
| chr12 | Oeu037980.1 | peroxidase 5-like | F:GO:0004601:peroxidase activity; P:GO:0006979:response to oxidative stress; F:GO:0020037:heme binding |
| chr12 | Oeu037981.1 | uncharacterized protein LOC111373264 | |

| | | | |
|---|---|---|---|
| chr12 | Oeu037982.1 | heavy metal-associated isoprenylated plant 9 | F:GO:0046872:metal ion binding |
| chr12 | Oeu037983.1 | 60S ribosomal protein L9 | F:GO:0003735:structural constituent of ribosome; C:GO:0005840:ribosome; P:GO:0006412:translation; F:GO:0019843:rRNA binding |
| chr12 | Oeu037984.1 | uncharacterized protein LOC105159007 | C:GO:0005576:extracellular region; P:GO:0060320:rejection of self pollen |
| chr12 | Oeu050214.1 | transcription factor MYB12-like | |
| chr12 | Oeu057647.1 | hypothetical protein H0E87_010871 | |
| chr12 | Oeu057648.1 | pentatricopeptide repeat-containing protein At2g03380, mitochondrial-like isoform X1 | F:GO:0005515:protein binding |
| chr12 | Oeu057649.1 | uncharacterized protein LOC111406154 isoform X1 | |
| chr12 | Oeu057651.1 | uncharacterized protein LOC111406155 | |
| chr12 | Oeu057652.1 | eukaryotic translation initiation factor 5A-2-like | F:GO:0003746:translation elongation factor activity; F:GO:0043022:ribosome binding; P:GO:0045901:positive regulation of translational elongation; P:GO:0045905:positive regulation of translational termination |
| scaffold1196 | Oeu005322.1 | ADP-ribosylation factor 1-like 2 isoform X1 | F:GO:0003924:GTPase activity; F:GO:0005525:GTP binding |
| scaffold1196 | Oeu005323.1 | ---NA--- | |
| scaffold1196 | Oeu005324.1 | serine/threonine-protein phosphatase 4 regulatory subunit 2 isoform X1 | F:GO:0019888:protein phosphatase regulator activity; C:GO:0030289:protein phosphatase 4 complex |
| scaffold1196 | Oeu005325.1 | importin-5-like | P:GO:0006606:protein import into nucleus |
| scaffold1196 | Oeu005327.1 | homeobox-leucine zipper protein ATHB-17-like | F:GO:0003677:DNA binding |
| scaffold1196 | Oeu005328.1 | uncharacterized protein LOC111385861 | |
| scaffold1264 | Oeu007174.1 | receptor kinase At4g00960 | |
| scaffold1264 | Oeu007175.1 | ---NA--- | |
| scaffold1264 | Oeu007176.1 | Hypothetical predicted protein | |
| scaffold1264 | Oeu007178.1 | phosphoglucomutase, cytoplasmic | P:GO:0005975:carbohydrate metabolic process; F:GO:0016868:intramolecular transferase activity, phosphotransferases |
| scaffold1264 | Oeu007179.1 | glucan endo-1,3-beta-glucosidase 3-like isoform X1 | F:GO:0004553:hydrolase activity, hydrolyzing O-glycosyl compounds; P:GO:0005975:carbohydrate metabolic process |
| scaffold1264 | Oeu007181.1 | probable peroxygenase 4 | |
| scaffold1264 | Oeu007182.1 | ubiquitin-conjugating enzyme E2 variant 1A-like | |
| scaffold1264 | Oeu007185.1 | uncharacterized protein LOC111394498 | |
| scaffold1264 | Oeu007186.1 | E3 ubiquitin- ligase RHA2A-like | |

| | | | |
|---|---|---|---|
| scaffold1264 | Oeu007187.1 | zinc finger CCCH domain-containing protein 15-like | F:GO:0046872:metal ion binding |
| scaffold1264 | Oeu007190.2 | lipid phosphate phosphatase 2-like | P:GO:0006644:phospholipid metabolic process; F:GO:0042577:lipid phosphatase activity |
| scaffold1264 | Oeu007191.1 | lipid phosphate phosphatase 2-like | P:GO:0006644:phospholipid metabolic process; F:GO:0042577:lipid phosphatase activity |
| scaffold1264 | Oeu007192.1 | uncharacterized protein LOC111374612 | F:GO:0005515:protein binding |
| scaffold393 | Oeu041861.2 | tRNA(adenine(34)) deaminase, chloroplastic-like | |
| scaffold393 | Oeu041862.2 | phosphate transporter PHO1 homolog 1-like isoform X1 | C:GO:0016021:integral component of membrane |
| scaffold393 | Oeu041863.1 | uncharacterized protein LOC111367484 | |
| scaffold393 | Oeu041866.1 | uncharacterized protein LOC111392799 | |
| scaffold393 | Oeu041867.1 | ATPase subunit 4 | C:GO:0000276:mitochondrial proton-transporting ATP synthase complex, coupling factor F(o); F:GO:0015078:proton transmembrane transporter activity; P:GO:0015986:ATP synthesis coupled proton transport |
| scaffold393 | Oeu041868.1 | ---NA--- | |
| scaffold393 | Oeu041869.1 | ---NA--- | |
| scaffold393 | Oeu041870.1 | protein BOBBER 2-like | |
| scaffold393 | Oeu041871.1 | axoneme-associated protein mst101(2)-like | |
| scaffold393 | Oeu041872.1 | ---NA--- | |
| scaffold393 | Oeu041874.1 | protein decapping 5-like | |
| scaffold393 | Oeu041875.1 | protein CROWDED NUCLEI 2-like | C:GO:0005634:nucleus; P:GO:0006997:nucleus organization |
| scaffold393 | Oeu041876.1 | Hypothetical predicted protein | |
| scaffold393 | Oeu041877.1 | Hypothetical predicted protein | |
| scaffold393 | Oeu041878.1 | Hypothetical predicted protein | |
| scaffold393 | Oeu041879.2 | SUPPRESSOR OF GAMMA RESPONSE 1-like | F:GO:0003677:DNA binding; F:GO:0003700:DNA-binding transcription factor activity; P:GO:0006355:regulation of transcription, DNA-templated |
| scaffold393 | Oeu041880.1 | ---NA--- | |
| scaffold393 | Oeu041881.1 | uncharacterized protein LOC111386005 | |
| scaffold393 | Oeu041882.1 | trafficking particle complex subunit 11 | |
| scaffold393 | Oeu041883.1 | transcription factor HHO3-like | F:GO:0003677:DNA binding; F:GO:0003700:DNA-binding transcription factor activity; P:GO:0006355:regulation of transcription, DNA-templated |

| | | | |
|---|---|---|---|
| scaffold393 | Oeu041885.1 | transcription factor HHO3-like | F:GO:0003677:DNA binding; F:GO:0003700:DNA-binding transcription factor activity; P:GO:0006355:regulation of transcription, DNA-templated |
| scaffold393 | Oeu041887.1 | Hypothetical predicted protein | |
| scaffold393 | Oeu041888.1 | ---NA--- | |
| scaffold393 | Oeu041891.1 | GDT1-like protein 4 isoform X1 | |
| scaffold393 | Oeu041892.2 | CTL-like protein DDB_G0274487 isoform X1 | F:GO:0022857:transmembrane transporter activity; P:GO:0055085:transmembrane transport |
| scaffold393 | Oeu041893.1 | uncharacterized protein LOC111387182 | |
| scaffold393 | Oeu041894.1 | serine/threonine-protein phosphatase 2A 65 kDa regulatory subunit A beta isoform-like | F:GO:0005515:protein binding |
| scaffold393 | Oeu041896.1 | heavy metal-associated isoprenylated plant protein 6-like | |
| scaffold393 | Oeu041897.1 | heat stress transcription factor C-1-like isoform X1 | F:GO:0003700:DNA-binding transcription factor activity; P:GO:0006355:regulation of transcription, DNA-templated; F:GO:0043565:sequence-specific DNA binding |
| scaffold393 | Oeu041898.1 | heat stress transcription factor C-1 | |
| scaffold393 | Oeu041901.1 | transcription factor MYB106-like | |
| scaffold393 | Oeu041902.1 | ---NA--- | |
| scaffold969 | Oeu064202.3 | hypothetical protein DKX38_015652 | F:GO:0005515:protein binding |
| scaffold969 | Oeu064204.1 | ---NA--- | |
| scaffold969 | Oeu064205.1 | vacuolar-sorting protein BRO1-like | F:GO:0005515:protein binding |
| scaffold969 | Oeu064206.1 | ATP synthase subunit beta, mitochondrial-like | C:GO:0000275:mitochondrial proton-transporting ATP synthase complex, catalytic sector F(1); F:GO:0005524:ATP binding; P:GO:0015986:ATP synthesis coupled proton transport; F:GO:0016887:ATP hydrolysis activity; F:GO:0046933:proton-transporting ATP synthase activity, rotational mechanism |
| scaffold969 | Oeu064207.2 | origin of replication complex subunit 4 isoform X4 | C:GO:0000808:origin recognition complex; F:GO:0003677:DNA binding; F:GO:0005524:ATP binding; C:GO:0005634:nucleus; P:GO:0006260:DNA replication; F:GO:0016887:ATP hydrolysis activity |
| scaffold969 | Oeu064208.1 | nuclear pore complex protein NUP155 isoform X1 | C:GO:0005643:nuclear pore; P:GO:0006913:nucleocytoplasmic transport; F:GO:0017056:structural constituent of nuclear pore |

Table S2. List of the 32 gene annotations in the chromosomal interval of the olive tree genome bounded by *P. angustifolia* loci strictly associated with SI. This list is completed by the 111 gene annotations in the eight scaffolds containing loci strongly or loosely associated with SI in *P. angustifolia*. The description field is taken from Unver *et al.* (2020) and is based on automated blast annotation. NA corresponds to predicted gene models with no hit.

| Genome location | Sequence ID | Description | GO annotation |
|---|---|---|---|
| chr18 | Oeu052727.1 | uncharacterized protein LOC111404303 | |
| chr18 | Oeu037727.1 | GATA transcription factor 5-like | P:GO:0006355:regulation of transcription, DNA-templated; F:GO:0008270:zinc ion binding; F:GO:0043565:sequence-specific DNA binding |
| chr18 | Oeu016310.2 | uncharacterized protein LOC111372111 isoform X1 | |
| chr18 | Oeu016307.4 | Hypothetical predicted protein | |
| chr18 | Oeu016306.2 | tubby-like F-box protein 5 isoform X1 | F:GO:0005515:protein binding |
| chr18 | Oeu037732.1 | uncharacterized protein LOC111367390 | |
| chr18 | Oeu037717.1 | protein SHI RELATED SEQUENCE 1-like isoform X1 | |
| chr18 | Oeu037714.1 | ---NA--- | |
| chr18 | Oeu052725.1 | CHAPERONE-LIKE PROTEIN OF POR1, chloroplastic | |
| chr18 | Oeu037735.1 | zinc finger MYM-type protein 1-like | |
| chr18 | Oeu037740.1 | uncharacterized protein LOC111373987 | |
| chr18 | Oeu052728.1 | uncharacterized protein LOC111372566 | |
| chr18 | Oeu037721.1 | hypothetical protein CRG98_024810 | F:GO:0008168:methyltransferase activity; C:GO:0016021:integral component of membrane; P:GO:0032259:methylation |
| chr18 | Oeu037725.1 | uncharacterized protein LOC111367386 | |
| chr18 | Oeu037716.1 | protein FAR1-RELATED SEQUENCE 4-like | |
| chr18 | Oeu037719.1 | ---NA--- | |
| chr18 | Oeu037736.3 | transmembrane and coiled-coil domain-containing protein 4 | |
| chr18 | Oeu037737.1 | ---NA--- | |
| chr18 | Oeu037738.1 | uncharacterized protein LOC111394886 | |
| chr18 | Oeu016311.1 | uncharacterized protein LOC111384618 | |
| chr18 | Oeu037724.1 | uncharacterized protein LOC111367386 | |
| chr18 | Oeu037733.1 | uncharacterized protein LOC111378408 | |
| chr18 | Oeu037720.1 | uncharacterized protein LOC111368283 | |
| chr18 | Oeu037730.1 | ---NA--- | |
| chr18 | Oeu037734.1 | ---NA--- | |
| chr18 | Oeu016314.1 | microtubule-associated TORTIFOLIA1 | C:GO:0005874:microtubule; F:GO:0008017:microtubule binding; C:GO:0045298:tubulin complex |
| chr18 | Oeu037731.1 | cytochrome P450 84A1 | F:GO:0004497:monooxygenase activity; F:GO:0005506:iron ion binding; F:GO:0016705:oxidoreductase activity, acting on paired donors, with incorporation or reduction of molecular oxygen; F:GO:0020037:heme binding |

| | | | |
|---|---|---|---|
| chr18 | Oeu037741.1 | ---NA--- | |
| chr18 | Oeu037739.1 | ---NA--- | F:GO:0003676:nucleic acid binding; F:GO:0008270:zinc ion binding |
| chr18 | Oeu052726.1 | ---NA--- | F:GO:0003677:DNA binding; F:GO:0005515:protein binding; F:GO:0046872:metal ion binding |
| chr18 | Oeu016308.1 | ---NA--- | F:GO:0004842:ubiquitin-protein transferase activity; P:GO:0016567:protein ubiquitination |
| chr18 | Oeu016312.1 | uncharacterized protein LOC111373662 | |
| scaffold1199 | Oeu005362.1 | uncharacterized protein LOC111366962 | |
| scaffold1199 | Oeu005363.1 | Hypothetical predicted protein | |
| scaffold1199 | Oeu005364.1 | protein indeterminate-domain 1-like isoform X1 | |
| scaffold1199 | Oeu005367.1 | deSI-like protein At4g17486 | F:GO:0008233:peptidase activity |
| scaffold1199 | Oeu005368.1 | peroxisomal membrane protein 11A | C:GO:0005779:integral component of peroxisomal membrane; P:GO:0016559:peroxisome fission |
| scaffold1199 | Oeu005369.2 | protein BRASSINAZOLE-RESISTANT 1-like | F:GO:0003700:DNA-binding transcription factor activity; P:GO:0006351:transcription, DNA-templated; P:GO:0009742:brassinosteroid mediated signaling pathway |
| scaffold1199 | Oeu005370.1 | NEDD8-conjugating enzyme Ubc12-like | F:GO:0005524:ATP binding; F:GO:0016740:transferase activity |
| scaffold1199 | Oeu005371.1 | 30S ribosomal protein S13, chloroplastic-like | F:GO:0003723:RNA binding; F:GO:0003735:structural constituent of ribosome; C:GO:0005840:ribosome; P:GO:0006412:translation |
| scaffold1199 | Oeu005372.1 | terminal ear1 homolog | F:GO:0003676:nucleic acid binding |
| scaffold1200 | Oeu005581.1 | transmembrane protein 87A-like | C:GO:0016021:integral component of membrane |
| scaffold1200 | Oeu005582.1 | exocyst complex component EXO70A1-like | C:GO:0000145:exocyst; P:GO:0006887:exocytosis |
| scaffold1200 | Oeu005583.1 | protein TIFY 11B-like | |
| scaffold1200 | Oeu005584.1 | putative B3 domain-containing protein Os03g0621600 | |
| scaffold1200 | Oeu005585.1 | zinc finger CCCH domain-containing 48-like isoform X2 | F:GO:0005515:protein binding; P:GO:0006364:rRNA processing; F:GO:0034511:U3 snoRNA binding; F:GO:0046872:metal ion binding |
| scaffold1200 | Oeu005587.1 | hypothetical protein CK203_000394 | |
| scaffold1200 | Oeu005588.1 | zinc finger CCCH domain-containing protein 62-like | F:GO:0003723:RNA binding; P:GO:0045892:negative regulation of transcription, DNA-templated; F:GO:0046872:metal ion binding |
| scaffold1200 | Oeu005589.1 | 25.3 kDa vesicle transport protein-like isoform X1 | F:GO:0005484:SNAP receptor activity; P:GO:0006888:endoplasmic reticulum to Golgi vesicle-mediated transport; P:GO:0006890:retrograde vesicle-mediated transport, Golgi to endoplasmic reticulum; C:GO:0016021:integral component of membrane |
| scaffold1200 | Oeu005590.1 | Hypothetical predicted protein | |
| scaffold1200 | Oeu005591.1 | ABC transporter B family member 19-like | F:GO:0005524:ATP binding; C:GO:0016021:integral component of membrane; P:GO:0055085:transmembrane transport; F:GO:0140359:ABC-type transporter activity |
| scaffold1287 | Oeu007808.1 | ---NA--- | |
| scaffold1287 | Oeu007810.1 | Hypothetical predicted protein | |
| scaffold1287 | Oeu007811.1 | Hypothetical predicted protein | |
| scaffold1287 | Oeu007813.1 | ---NA--- | |
| scaffold1287 | Oeu007814.1 | probable inactive poly [ADP-ribose] polymerase SRO3 | F:GO:0003950:NAD+ ADP-ribosyltransferase activity |

| | | | |
|---|---|---|---|
| scaffold1287 | Oeu007815.1 | KH domain-containing protein HEN4-like isoform X1 | F:GO:0003723:RNA binding |
| scaffold1579 | Oeu014557.1 | zinc finger BED domain-containing protein RICESLEEPER 2-like | |
| scaffold1579 | Oeu014558.1 | nitrate regulatory gene2 protein | |
| scaffold1579 | Oeu014559.1 | nitrate regulatory gene2 protein | |
| scaffold1579 | Oeu014560.1 | protein POLLENLESS 3-LIKE 2 | F:GO:0005515:protein binding |
| scaffold1579 | Oeu014561.2 | histidine kinase 1 isoform X2 | F:GO:0000155:phosphorelay sensor kinase activity; P:GO:0000160:phosphorelay signal transduction system; P:GO:0016310:phosphorylation |
| scaffold1579 | Oeu014562.1 | protein EARLY-RESPONSIVE TO DEHYDRATION 7, chloroplastic-like | |
| scaffold1579 | Oeu014563.1 | Hypothetical predicted protein | |
| scaffold1579 | Oeu014567.1 | alpha-mannosidase-like | F:GO:0004559:alpha-mannosidase activity; P:GO:0006013:mannose metabolic process; F:GO:0030246:carbohydrate binding |
| scaffold213 | Oeu024812.1 | uncharacterized protein LOC111367376 isoform X1 | P:GO:0007064:mitotic sister chromatid cohesion |
| scaffold213 | Oeu024815.1 | zinc finger BED domain-containing protein RICESLEEPER 1-like | F:GO:0003677:DNA binding; F:GO:0046983:protein dimerization activity |
| scaffold213 | Oeu024816.1 | Hypothetical predicted protein | |
| scaffold213 | Oeu024817.1 | uncharacterized protein LOC111380358 | |
| scaffold213 | Oeu024818.1 | blue copper -like | F:GO:0009055:electron transfer activity |
| scaffold213 | Oeu024819.1 | Hypothetical predicted protein | |
| scaffold213 | Oeu024820.1 | Hypothetical predicted protein | |
| scaffold213 | Oeu024822.1 | uncharacterized protein LOC120106791 | |
| scaffold213 | Oeu024824.1 | ribosomal protein S2 | F:GO:0003735:structural constituent of ribosome; P:GO:0006412:translation; C:GO:0009507:chloroplast; C:GO:0015935:small ribosomal subunit |
| scaffold213 | Oeu024825.1 | VQ motif-containing protein 9 | P:GO:1901001:negative regulation of response to salt stress |
| scaffold213 | Oeu024826.1 | Hypothetical predicted protein | |
| scaffold213 | Oeu024827.1 | uncharacterized protein LOC111380349 isoform X4 | |
| scaffold213 | Oeu024828.1 | AP-4 complex subunit mu-like | P:GO:0006886:intracellular protein transport; P:GO:0016192:vesicle-mediated transport; C:GO:0030131:clathrin adaptor complex |
| scaffold213 | Oeu024829.1 | uncharacterized protein LOC111406687 | |
| scaffold213 | Oeu024830.1 | ---NA--- | |
| scaffold213 | Oeu024832.1 | Hypothetical predicted protein | |
| scaffold269 | Oeu032218.1 | protein TsetseEP-like | |
| scaffold269 | Oeu032219.1 | uncharacterized protein LOC111383442 | |
| scaffold269 | Oeu032220.1 | DNA (cytosine-5)-methyltransferase 1B-like | F:GO:0003682:chromatin binding; F:GO:0003886:DNA (cytosine-5-)-methyltransferase activity; C:GO:0005634:nucleus; P:GO:0090116:C-5 methylation of cytosine |
| scaffold269 | Oeu032221.1 | ---NA--- | |

| | | | |
|---|---|---|---|
| scaffold269 | Oeu032222.1 | ---NA--- | |
| scaffold269 | Oeu032223.3 | uncharacterized protein LOC111383435 | |
| scaffold269 | Oeu032224.2 | membrin-11-like | |
| scaffold269 | Oeu032225.1 | hypothetical protein F0562_029638 | |
| scaffold269 | Oeu032226.1 | caltractin-like isoform X1 | F:GO:0005509:calcium ion binding |
| scaffold269 | Oeu032227.1 | uncharacterized protein LOC111398231 | |
| scaffold269 | Oeu032228.1 | uncharacterized protein LOC111394895 | F:GO:0008270:zinc ion binding |
| scaffold269 | Oeu032229.1 | zinc-finger homeodomain protein 5-like | |
| scaffold269 | Oeu032230.1 | protein SUPPRESSOR OF GENE SILENCING 3 | P:GO:0031047:gene silencing by RNA; P:GO:0051607:defense response to virus |
| scaffold269 | Oeu032231.1 | uncharacterized protein LOC111398231 | |
| scaffold269 | Oeu032236.1 | protein SUPPRESSOR OF GENE SILENCING 3-like | P:GO:0031047:gene silencing by RNA; P:GO:0051607:defense response to virus |
| scaffold327 | Oeu037602.1 | 60S ribosomal L34 | F:GO:0003735:structural constituent of ribosome; C:GO:0005840:ribosome; P:GO:0006412:translation |
| scaffold327 | Oeu037603.1 | uncharacterized protein LOC111373208 | |
| scaffold327 | Oeu037604.1 | uncharacterized protein LOC111373818 | |
| scaffold327 | Oeu037606.1 | Hypothetical predicted protein | |
| scaffold327 | Oeu037607.1 | probable carbohydrate esterase At4g34215 | |
| scaffold327 | Oeu037608.1 | probable carbohydrate esterase At4g34215 | |
| scaffold327 | Oeu037609.1 | Hypothetical predicted protein | |
| scaffold327 | Oeu037610.2 | BUD13 homolog | |
| scaffold327 | Oeu037611.1 | uncharacterized protein LOC111399181 | |
| scaffold327 | Oeu037612.1 | mannose-P-dolichol utilization defect 1 homolog 2-like | |
| scaffold327 | Oeu037613.1 | uncharacterized protein LOC111394498 | |
| scaffold327 | Oeu037614.1 | uncharacterized protein LOC111385761 | |
| scaffold327 | Oeu037615.1 | low-temperature-induced cysteine proteinase-like | P:GO:0006508:proteolysis; F:GO:0008234:cysteine-type peptidase activity |
| scaffold327 | Oeu037616.1 | Hypothetical predicted protein | |
| scaffold327 | Oeu037617.1 | ---NA--- | |
| scaffold327 | Oeu037618.1 | Hypothetical predicted protein | |
| scaffold327 | Oeu037619.1 | probable inactive receptor kinase At5g67200 | F:GO:0004672:protein kinase activity; F:GO:0005515:protein binding; F:GO:0005524:ATP binding; P:GO:0006468:protein phosphorylation |
| scaffold327 | Oeu037620.1 | uncharacterized protein LOC111385749 | |
| scaffold327 | Oeu037621.1 | kinesin-like protein KIN-4A isoform X1 | F:GO:0003777:microtubule motor activity; F:GO:0005524:ATP binding; P:GO:0007018:microtubule-based movement; F:GO:0008017:microtubule binding |
| scaffold327 | Oeu037623.1 | ethylene-responsive transcription factor ERF008-like | F:GO:0003677:DNA binding; F:GO:0003700:DNA-binding transcription factor activity; P:GO:0006355:regulation of transcription, DNA-templated |

| | | | |
|---|---|---|---|
| scaffold327 | Oeu037624.1 | Hypothetical predicted protein | |
| scaffold502 | Oeu047735.1 | purple acid phosphatase 2-like | F:GO:0003993:acid phosphatase activity; F:GO:0046872:metal ion binding |
| scaffold502 | Oeu047736.1 | Hypothetical predicted protein | |
| scaffold502 | Oeu047738.1 | WALLS ARE THIN 1 | C:GO:0016021:integral component of membrane; F:GO:0022857:transmembrane transporter activity |
| scaffold502 | Oeu047739.1 | unnamed protein product, partial | F:GO:0005198:structural molecule activity; F:GO:0005515:protein binding; P:GO:0006886:intracellular protein transport; P:GO:0016192:vesicle-mediated transport; C:GO:0030126:COPI vesicle coat |
| scaffold502 | Oeu047747.1 | B-box zinc finger 21-like | F:GO:0008270:zinc ion binding |
| scaffold502 | Oeu047748.1 | uncharacterized TPR repeat-containing protein At1g05150-like | F:GO:0005509:calcium ion binding; F:GO:0005515:protein binding |
| scaffold502 | Oeu047750.1 | probable 1-acylglycerol-3-phosphate O-acyltransferase | F:GO:0003824:catalytic activity |
| scaffold502 | Oeu047751.2 | hypothetical protein GIB67_019500 | F:GO:0015078:proton transmembrane transporter activity; C:GO:0016021:integral component of membrane; C:GO:0033179:proton-transporting V-type ATPase, V0 domain; P:GO:1902600:proton transmembrane transport |
| scaffold502 | Oeu047754.1 | importin beta-like SAD2 | |
| scaffold502 | Oeu047755.1 | cytokinin dehydrogenase 5-like | P:GO:0009690:cytokinin metabolic process; F:GO:0019139:cytokinin dehydrogenase activity; F:GO:0050660:flavin adenine dinucleotide binding |
| scaffold502 | Oeu047756.1 | Hypothetical predicted protein | |
| scaffold502 | Oeu047757.1 | Hypothetical predicted protein | |
| scaffold502 | Oeu047759.1 | probable ubiquitin-conjugating enzyme E2 18 | |
| scaffold502 | Oeu047760.1 | bZIP transcription factor 11-like | F:GO:0003700:DNA-binding transcription factor activity; P:GO:0006355:regulation of transcription, DNA-templated |
| scaffold502 | Oeu047761.1 | bZIP transcription factor 11-like | F:GO:0003700:DNA-binding transcription factor activity; P:GO:0006355:regulation of transcription, DNA-templated |
| scaffold502 | Oeu047764.1 | Hypothetical predicted protein | |
| scaffold502 | Oeu047765.2 | bifunctional nuclease 2-like | F:GO:0004518:nuclease activity |
| scaffold502 | Oeu047766.1 | ---NA--- | |
| scaffold502 | Oeu047767.2 | uncharacterized protein LOC111400469 | F:GO:0003676:nucleic acid binding; F:GO:0008270:zinc ion binding |
| scaffold502 | Oeu047769.1 | 50S ribosomal protein L31, chloroplastic-like | F:GO:0003735:structural constituent of ribosome; C:GO:0005840:ribosome; P:GO:0006412:translation |
| scaffold502 | Oeu047770.2 | zinc finger MYM-type protein 1-like | F:GO:0046983:protein dimerization activity |
| scaffold502 | Oeu047772.1 | putative pectinesterase 11 | F:GO:0030599:pectinesterase activity; P:GO:0042545:cell wall modification |
| scaffold502 | Oeu047773.1 | uncharacterized protein LOC111409242 isoform X1 | |
| scaffold502 | Oeu047774.1 | cell division homolog 2-1, chloroplastic-like | F:GO:0003924:GTPase activity; F:GO:0005525:GTP binding |
| scaffold502 | Oeu047775.1 | uncharacterized protein LOC111391588 | |
| scaffold502 | Oeu047778.1 | Erythronate-4-phosphate dehydrogenase family | |
